# Novel insights into joint estimations of demography, mutation rate, and selection using UV sex chromosomes

**DOI:** 10.1101/2021.03.30.437085

**Authors:** Sarah B. Carey, James H. Peniston, Adam C. Payton, Min Kim, Anna Lipzen, Diane Bauer, Kathleen Lail, Chris Daum, Kerrie Barry, Jerry Jenkins, Jane Grimwood, Jeremy Schmutz, Stuart F. McDaniel

## Abstract

A central goal in evolutionary genomics is to understand the processes that shape genetic variation in natural populations. In anisogamous species, these processes may generate asymmetries between genes transmitted through sperm or eggs. The unique inheritance of sex chromosomes facilitates studying such asymmetries, but in many systems sex-biased mutation, demography, and selection are confounded with suppressed recombination in only one sex (the W in females, or the Y in males). However, in a UV sex-determination system, both sex chromosomes are sex-specific and experience suppressed recombination. Here we built a spatially-structured simulation to examine the effects of population density and sex-ratio on female and male effective population size in haploids and compare the results to polymorphism data from whole-genome resequencing of the moss *Ceratodon purpureus*. In the parameter space we simulated, males nearly always had a lower effective population size than females. Using the *C. purpureus* resequencing data, we found the U and V have lower nucleotide diversity than the autosomal mean, and the V is much lower than the U, however, we found no parameter set in the model that explained both the U/V and U/autosome ratios we observed. We next used standard molecular evolutionary analyses to test for sex-biased mutation and selection. We found that males had a higher mutation rate but that natural selection shapes variation on the UV sex chromosomes. All together the moss system highlights how anisogamy alone can exert a profound influence on genome-wide patterns of molecular evolution.

## Introduction

Anisogamy, the condition in which genetic information is transmitted from one generation to the next through two different sized gametes, is widely shared among eukaryotes. The smaller gametes, typically called sperm, are abundant and motile, while the larger gametes, typically called eggs, are less abundant, better provisioned, and often sessile or retained on the parent. In species with two separate sexes, males produce sperm and females produce eggs, but of course many hermaphroditic species are also anisogamous. The asymmetry in gamete transmission means that the demography of genes transmitted through the smaller gamete can differ dramatically from the demography of genes transmitted through the larger gamete, even under neutral-equilibrium conditions (Charlesworth, 2009). For an extreme example, consider a population in which each egg-donor makes a single egg, but a single sperm-donor fertilizes all the eggs, a pattern which maximizes the variance in reproductive success for the sperm donor and dramatically reduces effective population size (*N_e_*) (Crow & Kimura, 1970; Sewell Wright, 1938). Anisogamy may therefore modulate the strength of selection on genes influencing transmission through sperm or eggs, potentially with major evolutionary consequences.

Most alleles are expressed in both females and males and therefore are transmitted through both egg and sperm. Thus, the effects of anisogamy on patterns of polymorphism will be averaged out across much of the genome. The sex chromosomes, however, are a major exception, because their patterns of inheritance are correlated with gametic sex and therefore record the history of sex-specific evolutionary processes (Caballero, 1995; Charlesworth, 2009; Kirkpatrick & Hall, 2004; Lenormand & Dutheil, 2005; Pool & Nielsen, 2007). In species with an XY sex-determination system, the Y chromosome is transmitted through males, meaning Y-chromosome polymorphism is shaped by transmission through sperm. However, no X homolog is transmitted exclusively through eggs because the X chromosome passes through both sexes. Similarly, ZW systems share these asymmetries in transmission, although in this case the W is female specific. In contrast, the UV sex chromosomes found in haploid systems with genetically-determined separate sexes have symmetrical transmission; the U is transmitted through eggs while the V is transmitted through sperm (Bachtrog et al., 2011; S. Carey, Kollar, & McDaniel, 2020). This inheritance pattern facilitates direct comparisons between female and male *N_e_* using homologous loci on the U and V.

Under the infinite-sites model, the equilibrium level of neutral variation depends only on the mutation rate and *N_e_* (Kimura, 1971). Male mutation rates may be higher than female rates, a finding that is often attributed to an increased number of cell divisions in the male germline (Hurst & Ellegren, 1998). It is unclear that this bias should apply to UV systems, which generally lack a distinction between the germline and soma. Even if the sexes have identical mutation rates, the levels of sex chromosome polymorphism are expected to be different than that for an autosome simply due to their mode of inheritance. For example, in diploid systems, each mated pair has three copies of an X or Z chromosome compared to four copies of each autosome and only one Y or W. Thus, the expected *N_e_* for an X or Z-linked locus is 3/4 of an autosome, while the expected *N_e_* for a Y or W-linked locus is 1/4 (when using discrete-generation approaches and assuming a Poisson offspring number distribution) (Charlesworth, 2001). In contrast, in haploid-dioecious systems, each mated pair has one U and one V for every two autosomes. Thus, both the U and the V chromosomes are expected to have 1/2 *N_e_* of an autosome, under similarly-restricted conditions (Avia et al., 2018; McDaniel, Neubig, Payton, Quatrano, & Cove, 2013). In UV systems, mitochondria and chloroplasts are maternally inherited (i.e., also sex specific) and expressed in the haploid stage, so they are also expected to have 1/2 *N_e_* of an autosome, while they are 1/4 *N_e_* in diploids (Sayres, 2018).

Numerous other non-random processes can also cause the *N_e_* of the sex chromosomes to deviate from the infinite-sites expectations. These processes may act in concert with, or independent of, the effects of anisogamy on *N_e_*. Sexual selection, for example, often generates greater variance in reproductive success in males, profoundly decreasing the *N_e_* of Y-linked loci, compared to X-linked loci or autosomes (Crow & Morton, 1955; Nunney, 1993). In discrete-generation models, a large excess in variance of male reproductive success over Poisson expectation causes ratios of sex chromosome and autosome polymorphism to approach extreme values (e.g., X/A=9/8, Y/A=1/8, and Y/X=1/9; (Caballero, 1995)). Sex-ratio biases can also drastically affect patterns of *N_e_*. For example, in an XY system, the ratio of X to autosomal diversity increases as the population becomes more female-biased, while the Y to autosome ratio decreases (Ellegren, 2009; Sayres, 2018). Additional demographic or life history factors may also drive species-specific variation in sex chromosome polymorphism. Age structure (Charlesworth, 2001) or geographic structure, in which males and females experience different migration patterns, can either moderate or exacerbate these biases (Goldberg & Rosenberg, 2015). Such processes and their effects on *N_e_* have attracted relatively little attention in bryophytes or other UV systems (but see (Bengtsson & Cronberg, 2009)).

Other forms of selection can also affect variation on sex chromosome and autosomes in different ways, potentially enhancing any asymmetries in *N_e_* among sex chromosomes. For example, the X chromosome is hemizygous in males, which can increase directional selection on male-beneficial recessive alleles and increase purifying selection on deleterious alleles (Charlesworth, Coyne, & Barton, 1987). Similarly, the male-specific region of the Y chromosome experiences suppressed recombination, meaning that linked selection drives patterns of polymorphism (Charlesworth & Charlesworth, 2000; J. M. Smith & Haigh, 1974). Polymorphism in mitochondrial and chloroplast DNA reflect female transmission and also exhibit suppressed recombination, but because they replicate independently in the cytoplasm they may experience unusual patterns of mutation or population size (D. R. Smith, 2015; Wolfe, Li, & Sharp, 1987). Thus, tests for sex-biased evolutionary processes in XY or ZW systems typically must rely upon comparisons among non-homologous loci that experience very different population genetic environments. In contrast, both the U and V experience suppressed recombination, meaning both the female and male-specific chromosomes are expected to experience an equivalent decrease in nucleotide diversity due to suppressed recombination (Avia et al., 2018; McDaniel, Neubig, et al., 2013).

Several sequenced UV sex chromosomes also maintain numerous homologs between the sexes, despite the suppressed recombination, providing abundant genetic data to study sex-specific variation (Ahmed et al., 2014; Bowman et al., 2017; S. B. Carey et al., 2020; Ferris et al., 2010). Sex chromosome degeneration in UV systems appears to be largely halted by strong purifying selection generated by haploid gene expression (S. B. Carey et al., 2020; Immler & Otto, 2015). One of the most remarkable examples is the moss *Ceratodon purpureus* in which the U and V sex chromosomes comprise ~30% of the ~360 megabase (Mb) genome for females and males, respectively (S. B. Carey et al., 2020). The U and V also contain ~12% of the organism’s gene content, providing numerous U-V orthologs to study differences in their molecular evolution (S. B. Carey et al., 2020). Nucleotide diversity data from a small number of U and V-linked introns in *C. purpureus* suggested that female and male-transmitted loci harbored similar amounts of genetic diversity, and both sexes showed indistinguishable patterns of population differentiation, suggesting that female and male spores may have equal probabilities of dispersing among populations (McDaniel, Neubig, et al., 2013). However, these data were insufficient to test for modest differences between female and male *N_e_* or mutation rate.

The life cycle of *C. purpureus* is like that of many dioecious species with UV sex-determination. Sexual reproduction typically occurs annually (Crum & Anderson, 1981; A. J. Shaw & Gaughan, 1993). Haploid males release V-carrying motile sperm, which either swim or are transported by microarthropods to egg-bearing females (Cronberg, Natcheva, & Hedlund, 2006; Rosenstiel, Shortlidge, Melnychenko, Pankow, & Eppley, 2012; Shortlidge et al., 2020), potentially meters away (Glime & Bisang, 2017). Haploid females make several identical U-carrying eggs, each enclosed in an archegonium. Although multiple eggs may be fertilized, typically only one embryo (i.e., sporophyte) develops. At maturity a sporophyte makes thousands of viable spores (Norrell, Jones, Payton, & McDaniel, 2014; A. J. Shaw & Gaughan, 1993; Shortlidge et al., 2020), most of which fall near the parent sporophyte, but some are captured by air currents and travel great distances (Biersma et al., 2020; McDaniel & Shaw, 2005).

Demographically-informed expectations that specifically incorporate anisogamy are necessary to fully understand the role of sex-specific evolutionary forces shaping patterns of polymorphism in dioecious species. In principle, a single male could fertilize many nearby females, an inference supported by field observations and experiments (Johnson & Shaw, 2016; Shortlidge et al., 2020), which increases variance in male reproductive success. Many bryophyte populations, including mosses, have an apparent female-biased sex ratio, due to sex-biased differences in clonal growth rates, differences in mortality, or differences in the number of fertile individuals during any given episode of reproduction, exacerbating this effect (Baughman, Payton, Paasch, Fisher, & McDaniel, 2017; Bisang, Ehrlén, & Hedenäs, 2019; Bisang & Hedenäs, 2005; Eppley et al., 2018; Norrell et al., 2014). The likelihood that a male sires offspring with multiple genetically-distinct females depends upon the spatial distribution of female and male genotypes, which in turn depends upon the recruitment of migrants and the clonal spread of the constituents of the population. Populations of *C. purpureus*, which are common in disturbed sites in temperate regions of all continents (Crum & Anderson, 1981), grow in dense patches with many distinct genotypes in close proximity (McDaniel & Shaw, 2005) and have a highly-variable sex ratio (Eppley et al., 2018). Here we built a simulation parameterized using life-history data from *C. purpureus* to evaluate the effect of demographic variables (population density and sex ratio) on female and male *N_e_*. We then compared the simulated data to patterns of polymorphism in genome-wide resequence data from *C. purpureus* to evaluate if the empirical observations could be explained by demographic processes alone. Our results highlight the importance of considering the joint effects of demography, selection, and mutation-rate variation in interpreting patterns of nucleotide polymorphism.

## Materials and Methods

In this manuscript, we use the term ‘female’ to describe individuals that inherit XX, ZW, or U chromosome(s) and produce eggs and we use the term ‘male’ to describe individuals that inherit XY, ZZ, or V chromosome(s) and produce sperm. We use this designation because it captures key aspects of transmission genetics, but we acknowledge that karyotypic sex does not always align with gametic sex, so this definition misses important components of diversity within a population or generation.

### Life history simulations

To generate demographically-informed estimates for *N_e_* of the U and V sex chromosomes (*N_eU_* and *N_eV_*, respectively), we constructed a spatially-explicit simulation. We made several assumptions based on the life cycle of *C. purpureus*, namely that every individual was either male or female (i.e., there were no hermaphroditic individuals), and its sex was genetically determined; that all reproduction occurs sexually; that mating occurs once per generation and only occurred between adjacent males and females; and that each female mated once per generation, but that a male could mate with any adjacent female (i.e., up to eight females in the grid used in the simulations) during a bout of mating. We made a few simplifying assumptions, namely that all individuals were capable of sexual reproduction; that there was no mate preference; and that each mating event within a generation resulted in the same number of offspring. To keep population size constant, we adjusted the fecundity so that the mean number of offspring per individual was two. Therefore, the fecundity *F* was given by *F* = 2/*r*, where *r* is the mean number of females that each male mated with that generation.

Relaxing these assumptions, which are clearly violated in nature (e.g., clonal growth is frequent) can increase the variance in reproductive success in either sex, but in general should not qualitatively affect the results.

For each run of the simulation (equivalent to one generation of mating), individuals were randomly placed onto a 100 × 100 cell grid. Females mated in a random order. Each female searched the eight adjacent cells to it for males to mate with. If there was more than one adjacent male, the female randomly selected one of them to mate with. If no males were in the cells adjacent to the female, that female did not mate. An example of the population following one run of the simulation can be seen in Figure S1. All simulations were run in R (3.5.1; (R Core Team, 2013) and plotted using the packages reshape2 (v1.4.3; (Wickham, 2007, 2012) and ggplot2 (v3.2.1; (Wickham, 2016)).

For each run of the simulation, we recorded the total number of offspring per each male and female (females could only have 0 or *F* offspring) and calculated the variance in reproductive success for each sex. The variances in reproductive success were then used to calculate the *N_e_* of the U (females) and V (males) chromosomes, using the following equations (1), respectively

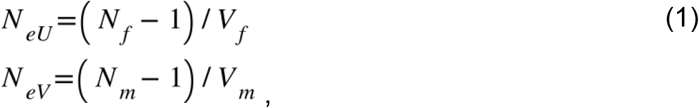

where *N_f_* and *N_m_* are the census population sizes and *V_f_* and *V_m_* are the variances in reproductive success of females and males, respectively. The *N_e_* of an autosome is calculated using the equation 2 (Crow & Kimura, 1970)

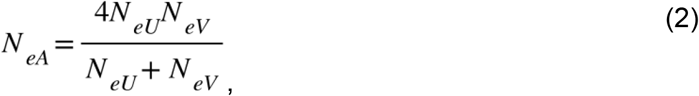

(derivations of equations 1 and 2 can be found in the Supplementary Appendix, Part 1).

To evaluate a variety of demographic scenarios, we ran simulations for a range of population densities and sex ratios. To vary density, we varied the population size while always keeping the arena of a fixed size of 100 × 100 cells. For each parameter set, we generated 100 runs of the simulation.

### Generating resequence data to test the model

We generated U-linked, V-linked, and autosomal polymorphism data from 23 *C. purpureus* isolates collected from nine locations (Figure 1; Table S1). To start these lines, sporophytes were surface sterilized, and a single germinated spore was isolated following (Norrell et al., 2014). DNA was extracted using a modified CTAB protocol following (Norrell et al., 2014). Plate-based DNA library preparation for Illumina sequencing was performed on the PerkinElmer Sciclone NGS robotic liquid handling system using Kapa Biosystems library preparation kit. Two hundred nanograms of sample DNA was sheared to 500 base pairs (bp) using a Covaris LE220 focused-ultrasonicator. The sheared DNA fragments were size selected by double-SPRI and then the selected fragments were end-repaired, A-tailed, and ligated with Illumina compatible sequencing adaptors from IDT containing a unique molecular index barcode for each sample library. The prepared sample libraries were quantified using KAPA Biosystem’s next-generation sequencing library qPCR kit and run on a Roche LightCycler 480 real-time PCR instrument. The quantified sample libraries were then multiplexed into pools and the pools were then prepared for sequencing on the Illumina HiSeq sequencing platform utilizing a TruSeq paired-end cluster kit v3 and Illumina’s cBot instrument to generate clustered flow cells for sequencing. Sequencing of the flow cells was performed on the Illumina HiSeq2000 sequencer using Illumina TruSeq SBS v3 sequencing kits, following a 2×150 indexed high-output run recipe. A subset of the libraries was also prepared using v4 chemistry and sequenced on a HiSeq2500 (see Table S1).

**Figure 1.**
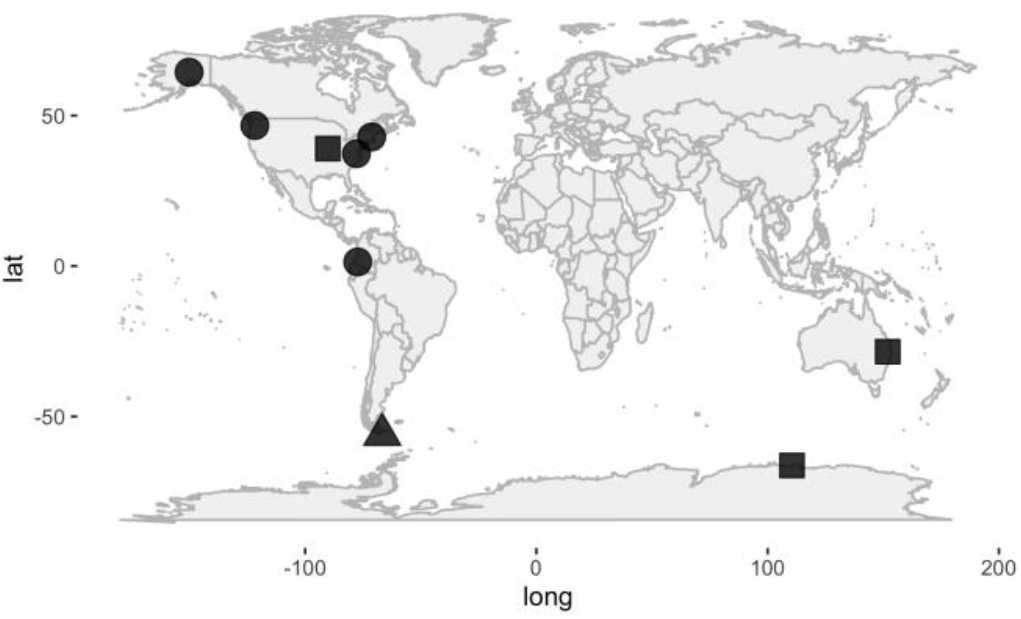
Sampling localities for the 23 *Ceratodon purpureus* isolates. All isolates from the nine localities were used for SNP calling. Some isolates were not used for downstream analyses because all isolates from the locality were female (indicated with squares). Isolates indicated with circles were used for the population genetics analyses and the isolates indicated with a triangle were used as the outgroup for the McDonald Kreitman Test.

From the raw reads, we removed artifact sequences, reads containing N bases, low-quality reads, DNA spike-in sequences, and PhiX control sequences. We split paired-end reads into forward and reverse using an in-house script from (S. B. Carey et al., 2020). We removed Illumina adapters and further filtered for quality using Trimmomatic (v0.36; (Bolger, Lohse, & Usadel, 2014)) using leading and trailing values of three, window size of 10, quality score of 30, and minimum length of 40. We visually assessed the quality of the remaining reads using fastqc (v0.11.4; (Andrews, 2010)).

We determined the karyotypic sex of these isolates by mapping reads using HISAT2 (v2.1.0; (Kim, Langmead, & Salzberg, 2015)) to the *Ceratodon purpureus* v1.0 genome using the R40 isolate (autosomes and V sex chromosome) concatenated with the GG1 U sex chromosome (S. B. Carey et al., 2020). We converted the resulting SAM files to sorted BAMs and indexed using SAMtools (v1.9; (H. Li et al., 2009)). Using IGV (v2.15.0; (Robinson et al.,2011)), we visually assessed sex by determining to which sex chromosome the reads mapped at the oldest locus known to be sex-linked in mosses (CepurR40.VG235300 and CepurGG1.UG071900 from (S. B. Carey et al., 2020)) and haphazardly scanning along the sex chromosomes.

To map the reads for downstream molecular evolutionary analyses, we used the genome reference described above, but also included R40’s chloroplast assembly. The R40 chloroplast was assembled using NOVOPlasty v2.6.7 (Dierckxsens, Mardulyn, & Smits, 2017) from existing Illumina data deposited in the NCBI BioProject PRJNA258984 from (S. B. Carey et al., 2020). Due to the low divergence between much of the U and V sex chromosomes (S. B. Carey et al., 2020; McDaniel, Neubig, et al., 2013), to ensure isolates mapped to the correct sex chromosome we hard masked the U for males and V and for females using BEDTools *maskfasta* (v2.27.1; (Quinlan & Hall, 2010), following (Olney, Brotman, Andrews, Valverde-Vesling, & Wilson, 2020)). Previous analyses found *C. purpureus* was highly polymorphic (McDaniel, van Baren, Jones, Payton, & Quatrano, 2013), so we used two mappers, BWA-MEM (v0.7.17; (H. Li, 2013)) and NGM (v0.5.5; (Sedlazeck, Rescheneder, & von Haeseler, 2013)), as they handle divergence differently, and ran analyses on these separately. We added read groups to the SAM files using Picard Tools (v2.19.1; http://broadinstitute.github.io/picard) *AddOrReplaceReadGroups* and converted them to sorted BAMs using SAMtools (v1.9; (H. Li et al., 2009)).

We called variants on all BAMs together using BCFtools (v1.9; (H. Li, 2011))*mpileup* and *call* using a ploidy of one. The resulting VCF file was filtered using BCFtools *filter* by excluding variants with a Phred-based quality score of the alternate base (QUAL) <30, combined depth across samples (DP) <10, and mapping quality (MQ) <30, where these filters had to be met in at least one sample (&&). We subset the VCFs using *view* to have females for the U, males for the V, and both sexes for the chloroplast and autosomal analyses, excluding isolates from localities where both sexes were not present (Figure 1; Table S1). The VCFs were finally filtered to remove variants with >20% missing data.

### Population genetic analyses

To examine patterns of nucleotide diversity, we calculated Wu and Watterson’s theta (*θ*; (Watterson, 1975)), which is based on the number of segregating sites in the population (*S*), and Nei and Li’s Pi (*π*), which is based on the average number of pairwise differences (Nei & Li, 1979), with both calculated per site (*N*_Localities_=5, *N*_Males_=8, *N*_Females_=8). We calculated Tajima’s D (Tajima, 1989) for the autosomes and sex chromosomes to test whether the mutation-frequency spectrum differed between these genomic regions, where a negative D suggests an excess of rare alleles indicating a recent selective sweep or expansion after a bottleneck, and a positive D suggests a lack of rare alleles indicating balancing selection or population contraction (*N*_Localities_=5, *N*_Males_=8, *N*_Females_=8). To examine differences in gene flow between autosomes and sex chromosomes we calculated *F*_ST_ (Sewall Wright, 1949), comparing the isolates from Alaska and Portland (*N*_Males_=4, *N*_Females_=4) to those from Durham and Storrs (*N*_Males_=3, *N*_females_=3) (Figure 1; Table S1). For each of these metrics we did sliding-window analyses using a window size of 100,000 and jump of 10,000 and plotted these with a loess correction span of 0.03 using karyoploteR (v1.8.8; (Gel & Serra, 2017)). We excluded the chloroplast, however, because the contig we analyzed was barely larger than the windows (105,555 bp). We generated 95% confidence intervals (CI) by bootstrapping 1,000 of the sliding windows with replacement and tested for differences between the autosomes and sex chromosomes using the Mann-Whitney U test with a Benjamini and Hochberg correction for multiple tests (Benjamini & Hochberg, 1995; McKnight & Najab, 2010).

To test for adaptive evolution in autosomal and sex-linked genes we first calculated the McDonald Krietman (MK) test (McDonald & Kreitman, 1991). The MK test compares the ratio of non-synonymous polymorphisms (*Pn*) to synonymous polymorphisms (*Ps*) to the ratio of non-synonymous divergence (*Dn*) to synonymous divergence (*Ds*), where under neutrality these two ratios are expected to be equal (i.e., (*Dn*/*Ds*) = (*Pn*/*Ps*)). Several phylogenetic analyses showed the Chilean populations were isolated from northern hemisphere populations, potentially representing a new species (Biersma et al., 2020; McDaniel & Shaw, 2005), so we used the female and male Chile isolates as the outgroup for the MK test (*N*_Localities_=6, *N*_Males_=9, *N*_Females_=9). We evaluated the significance of deviations from neutrality using Fisher’s exact test (Fisher, 1922). Finally, we calculated the Direction of Selection (DoS) test (Stoletzki & Eyre-Walker, 2011), using the equation

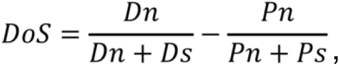

where a DoS < 0 indicates relaxed purifying selection and DoS > 0 indicates positive selection.

To determine if the mutation rate differed between the U and V chromosomes, using PAML (Yang, 2007) we calculated synonymous (*dS*) and nonsynonymous (*dN*) changes on branches of 330 one-to-one orthologs of the R40 and GG1 genome isolates. The gene trees used and in-depth details of running PAML were previously reported in (S. B. Carey et al.,2020). We tested the difference in *dN* and *dS* between the U and V-linked orthologs, using the Mann-Whitney U test (Benjamini & Hochberg, 1995; McKnight & Najab, 2010), removing one V-linked gene with *dS*>10. All population genetic analyses were done in R version 3.5.3 (R Core Team, 2013) using PopGenome (v2.7.1; (Pfeifer, Wittelsbürger, Ramos-Onsins, & Lercher, 2014)) and plotted using ggplot2 (v3.3.1; (Wickham, 2016)), unless otherwise stated.

### Comparing empirical data to simulations

The results of the simulations provide demographically-informed expectations for levels of polymorphism for various population densities and sex ratios. To compare our resequence data to the life history simulations, we calculated the variation in reproductive success of both males and females that would be necessary to explain our data in the absence of other processes (e.g., selection or migration). We then compared the ratios of variation in reproductive success to the results of our simulations to see if population density or sex ratio could explain the nucleotide diversity patterns we observed. To calculate the variation in reproductive success needed to explain our results, we used the result that, in haploid populations, *θ* = 2*N_e_μ*, where *μ* is the mutation rate, (derivation provided in the Supplementary Appendix, Part 2). From this it follows that, given equation 1, *θ* for the U and V chromosomes are respectively given by

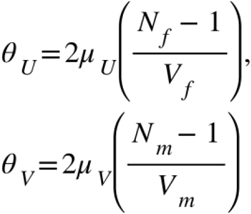

where *μ_U_* and *μ_V_* are the mutation rates on the U and V chromosomes, respectively. These equations can be rewritten in the form

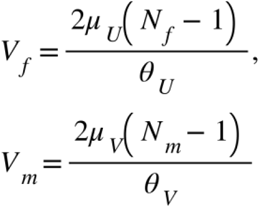

Similarly, it can be shown that

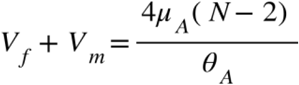

where *θ_A_* is the Wu and Watterson estimator for an autosome, *μ_A_* is the autosomal mutation rate, and *N* is the census population size assuming that the sex ratio is equal (a different equation is needed for unequal sex ratios).

From our resequence data, we estimated the ratio between *θ* for the V and U chromosomes, *θ_V_/θ_U_*, which we used to solve for the ratio *α* = *V_m_/V_f_* to determine how different the variances in reproductive success would need to be to explain the results, such that (7)

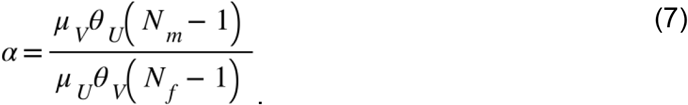

Similarly, given the ratio *θ_V_/θ_A_*, we can solve for the following ratio

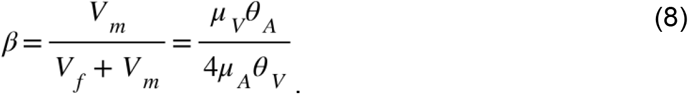

This solution for *β* does not hold for unequal sex ratios, however. Therefore, for unequal sex ratios we evaluated the ratio

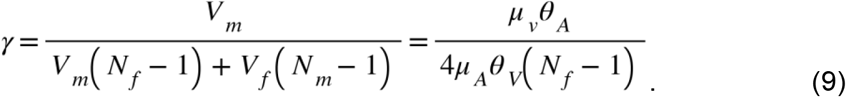

For calculations from our empirical data, we assume that *N* = 400,000, which is consistent with the *N_e_* calculated in *C. purpureus* (M. Nieto-Lugilde, *personal communication*).

## Results

### Life history simulations

To develop a demographically-informed model for sex-specific patterns of polymorphism, we used simulations to calculate *N_e_* for males and females at a range of population densities and sex-ratios. As the population density increased from 10–50% occupancy, with an equal number of males and females, *N_eA_, N_eU_*, and *N_eV_* increased, although not equivalently (Figure 2A). At the lowest densities, the *N_eA_* exceeded that of the U, and the V was lower still. All three *N_e_* values were well below the census population size, because relatively few females were adjacent to males, and therefore few individuals reproduced creating high variance in reproductive success. At ~25% occupancy, *N_eA_* was roughly equivalent to that of the U chromosome, while *N_eV_* was approximately half of the autosomal value. That is, variance in male reproductive success increases with increasing population density. At higher densities, the autosomal and V-chromosome *N_e_* increased approximately linearly, while *N_eU_* increased approximately exponentially, and at high densities the effective population of the U chromosome can exceed that of autosomes and even the census population size (Figure 2A). This seemingly counterintuitive finding arises from the fact that, at high population densities, the autosomal diversity is passed through relatively few males, while the U-linked variation is passed exclusively through the females which have very low variation in reproductive success.

**Figure 2.**
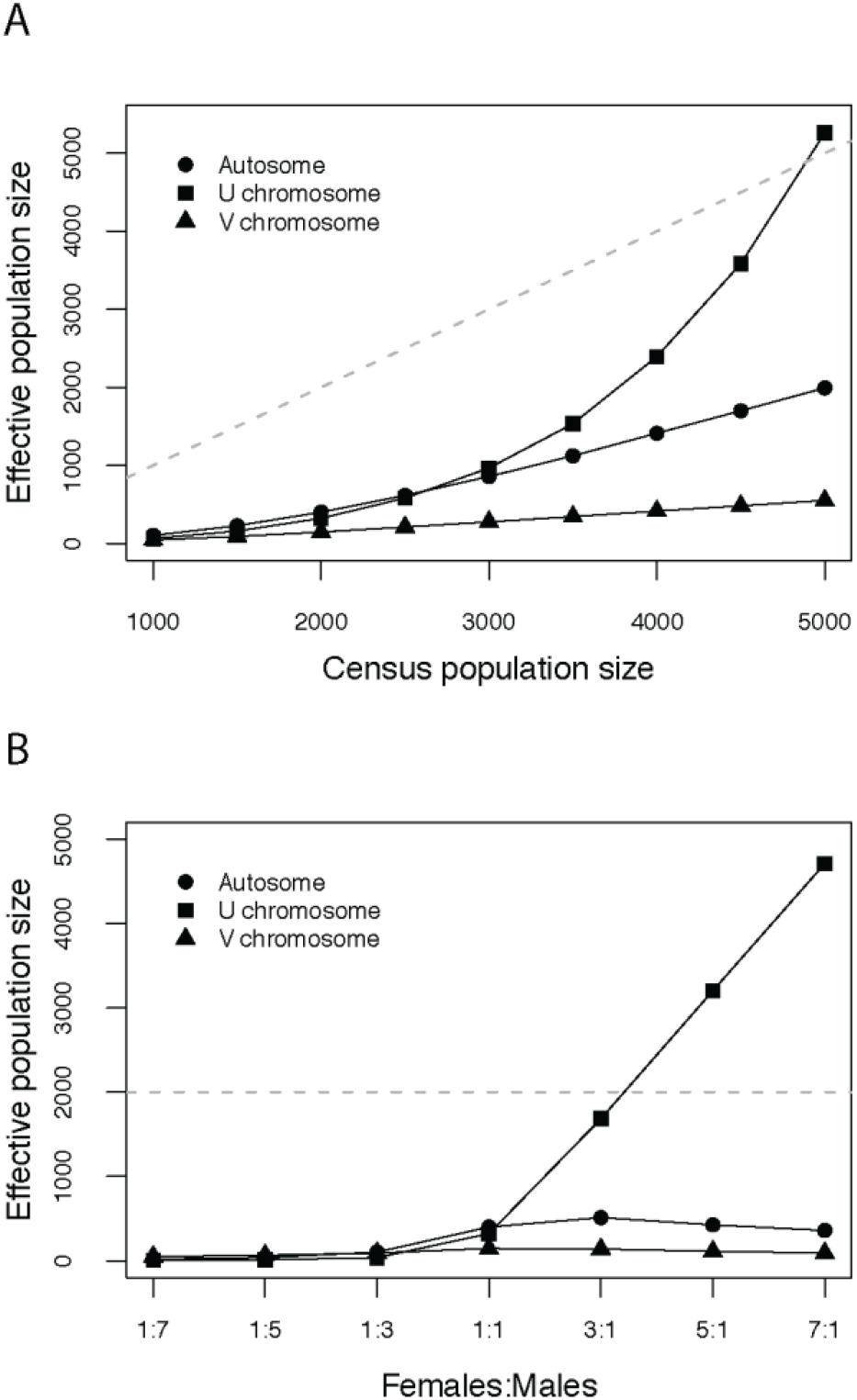
Effective population size of U and V sex chromosomes and autosomes calculated from spatially-explicit simulations. A) The effect of population density on effective population size (*N_e_*). In all simulations, there were 1000 available sites so density is census size divided by 1000. The dashed gray line denotes the one to one line at which the *N_e_* equals the census population size. B) The effect of sex ratio on *N_e_*. Dashed gray line shows the census population size of 2000. Each point is the mean value given by 100 runs of the simulation.

We also simulated the effect of variation in sex ratio on *N_e_* of the U, V, and autosomes. We explored the effects of sex-ratio variation at multiple population densities, but because the trends were homogeneous we present only the results at a density of 20%. At this population density, with an even sex ratio, the results most closely match the infinite-sites expectations. At even modest male-biased sex ratios, the *N_e_* of the U, V, and autosomes were very low (Figure 2B). Even when males outnumbered females, relatively few males contributed to reproduction. As the sex ratio became more female biased, in contrast, *N_eU_* increased dramatically (Figure 2B). The *N_eA_* increased slightly with a modest female bias, but at more dramatic female biases the *N_eA_* decreased slightly.

These models demonstrate that under reasonable demographic conditions, the infinite-sites expectations that the U and V each should have half the *N_e_* of an autosome are met only at low population densities. Moreover, the V can have lower *N_e_* than the U and autosomes due to a greater variance in reproductive success due to sex differences in life history alone (i.e., without selection or female mate choice), a pattern exacerbated by a sex-ratio bias toward either males or females.

### Patterns of polymorphism in *C. purpureus*

To empirically test our model of *N_e_* in a species with UV sex chromosomes, we generated whole-genome resequence data for 23 isolates of *C. purpureus* (Figure 1; Table S1). We found across isolates on average 80.87% of reads mapped with BWA and 81.64% with NGM and our average coverage is ~28.5x (Table S1). We found 21,907,382 SNPs using BWA and after filtering, for downstream analyses, we had 17,510,525 total SNPs, with 3,117,274 on the U, 2,372,026 on the V, and 12,021,225 on the autosomes and chloroplast. Using NGM we found 19,395,846 SNPs, and after filtering we had 11,580,361 SNPs on the autosomes and chloroplast, 2,397,292 on the U, and 1,660,989 on the V. Below for simplicity we discuss the remaining results from using the BWA mapper, although the summary statistics were similar with NGM and we report these in Table S2.

We found Wu and Watterson’s theta (*θ*) across the 12 autosomes (*θ_A_*) was on average 0.00983 (CI=0.00960-0.01025; Table 1). Given the relationship between *N_e_* and *θ* (i.e., *θ*=2*N_e_μ*), the U (*θ_U_*) and V (*θ_V_*) sex chromosomes and the chloroplast (*θ_C_*) should be 1/2 *θ_A_*, under neutral processes. However, we find *θ_U_*=0.00339 (CI=0.00329-0.00346), *θ_V_*=0.00241 (CI=0.00235-0.00248), and *θ_C_*=0.00015 (Table 1). Thus, the ratios for U/A =~1/3, U/A=1/4, and C/A =~1/40, rather than 1/2 for any of these chromosomes and V/U=~2/3 rather than 1. We found the same pattern using Nei and Li’s Pi (*π*) with *θ_A_*=0.00946 (CI=0.00916-0.00983), *π_U_*=0.00331 (CI=0.00323-0.0034), *π_V_*=0.00219 (CI=0.00213-0.00225), and *λ_C_*=0.00018 (Table 1). For Tajima’s D we found the autosomes on average were negative (−0.219; CI=(−0.325)-(−0.253)), as were both sex chromosomes (U −0.43, CI=(−0.415)-(−0.378); V −0.82, CI=(−0.826)-(−0.794)) and the chloroplast was positive (0.87) (Table 1). For *F*_ST_ between populations, the autosomes were on average 0.202 (CI=0.19-0.205), chloroplast 0.47, U 0.372 (CI=0.370-0.379), and V 0.375 (CI=0.376-0.385) (Table 1). We calculated sliding windows for these metrics, which show ample variation on the autosomes, but the sex chromosomes are homogenous (Figure 3). For all metrics, we found the autosomes, U, and V to be significantly different from each other (Mann Whitney U, p<0.00001).

**Figure 3.**
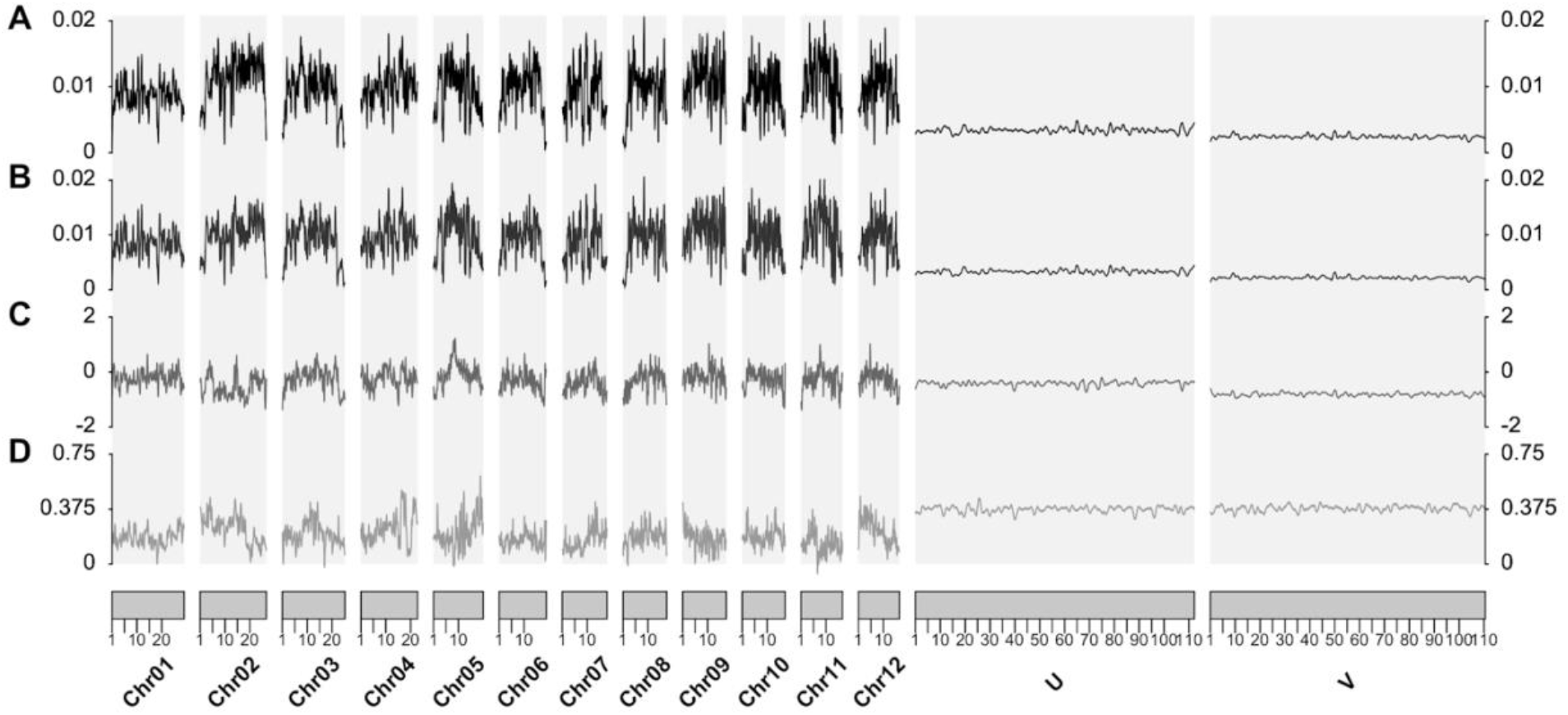
Sliding-window plots of nucleotide polymorphism summary statistics. Sliding windows for A) Wu and Watterson’s theta (*θ*), B) Pi (*π*), C) Tajima’s D, and D) *F*_ST_ were calculated using 100,000 bp windows with a 10,000 bp jump per chromosome. Lines were plotted using a loess correction with a span of 0.03. Note the different metrics have different y-axes, where *θ* and *π* range between 0 and 0.02, Tajima’s D ranges between −2 and 2, and *F*_ST_ ranges from 0 to 0.75.

**Table 1.** Population genetic analyses by chromosome. Segregating sites (*S*); Wu and Watterson’s theta (*θ*) Pi (*π*).

| Chromosome | Total sites | *S* | $\theta$ | $\pi$ | Tajima's D | *F<sub>ST</sub>* |
| --- | --- | --- | --- | --- | --- | --- |
| 1 | 29001003 | 829016 | 0.00868 | 0.00843 | -0.184 | 0.184 |
| 2 | 26629683 | 998325 | 0.01141 | 0.01015 | -0.534 | 0.247 |
| 3 | 25160467 | 770120 | 0.00929 | 0.00906 | -0.167 | 0.222 |
| 4 | 22785024 | 748029 | 0.00997 | 0.00965 | -0.197 | 0.268 |
| 5 | 19969424 | 641039 | 0.00975 | 0.00998 | 0.047 | 0.211 |
| 6 | 18980603 | 610384 | 0.00976 | 0.0092 | -0.312 | 0.168 |
| 7 | 17972677 | 533845 | 0.00902 | 0.00835 | -0.385 | 0.175 |
| 8 | 17567963 | 553075 | 0.00956 | 0.0092 | -0.221 | 0.202 |
| 9 | 17527894 | 624924 | 0.01087 | 0.0107 | -0.108 | 0.18 |
| 10 | 17229405 | 521827 | 0.00919 | 0.00891 | -0.192 | 0.195 |
| 11 | 16661191 | 593265 | 0.01082 | 0.01053 | -0.173 | 0.139 |
| 12 | 16459275 | 522515 | 0.00965 | 0.00931 | -0.207 | 0.236 |
| Chloroplast | 105555 | 51 | 0.00015 | 0.00018 | 0.868 | 0.471 |
| U | 112179120 | 982232 | 0.00339 | 0.00331 | -0.431 | 0.372 |
| V | 110524308 | 689145 | 0.00241 | 0.00219 | -0.818 | 0.375 |

We calculated the MK test on all 34,458 genes and found 280 had significant fixed amino acid changes relative to polymorphic changes based on Fisher’s exact test at p<0.05 and 606 at p<0.1 (Table 2, S3). Using the DoS test, we found for autosomes that 151 genes were less than one (at p<0.05; 338 at p<0.1) and 120 greater than one (p<0.05; 250 at p<0.1). For the U-linked genes, we found four genes were less than one and three greater than (at p<0.05; six at p<0.1). For V-linked genes we found two significant genes at p<0.05, with both less than one (5 at p<0.1) and two greater than one at p<0.1 (Figure 4A).

**Figure 4.**
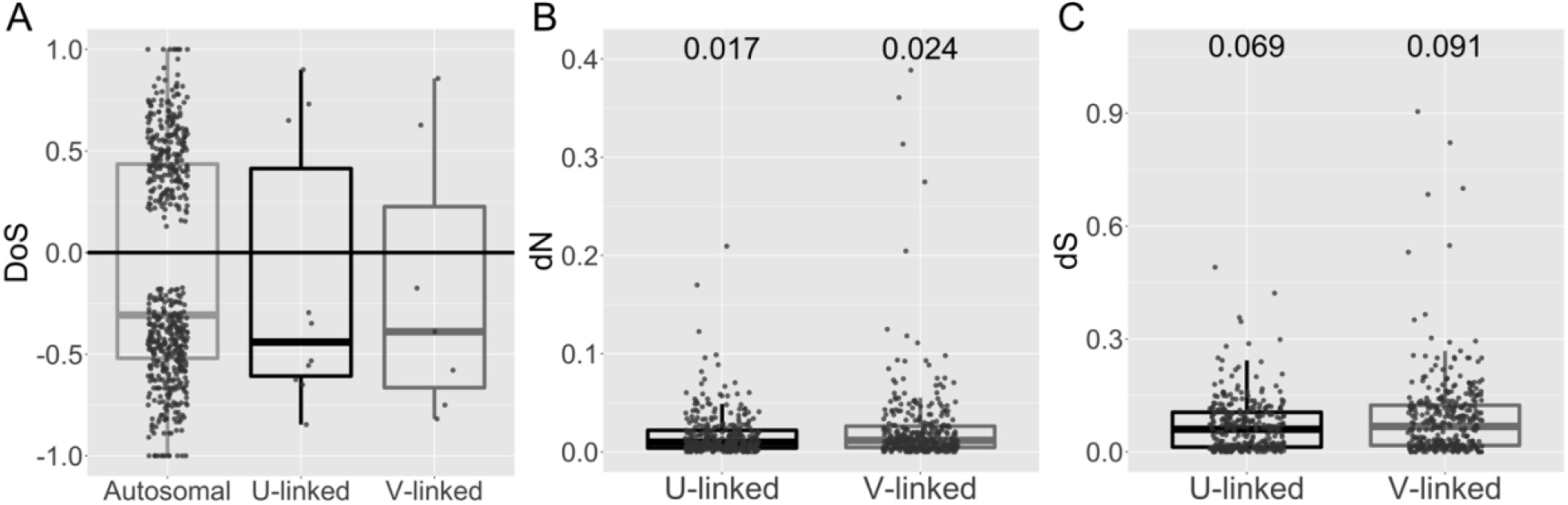
Measures of protein evolution. A) Direction of Selection (DoS) test for autosomal and sex-linked genes that were significant in the MK test at p<0.1. B) nonsynonymous mutation rate (*dN*) and C) synonymous mutation rate (*dS*) of one-to-one orthologous U and V-linked genes. Numbers on top show the mean values.

**Table 2.**
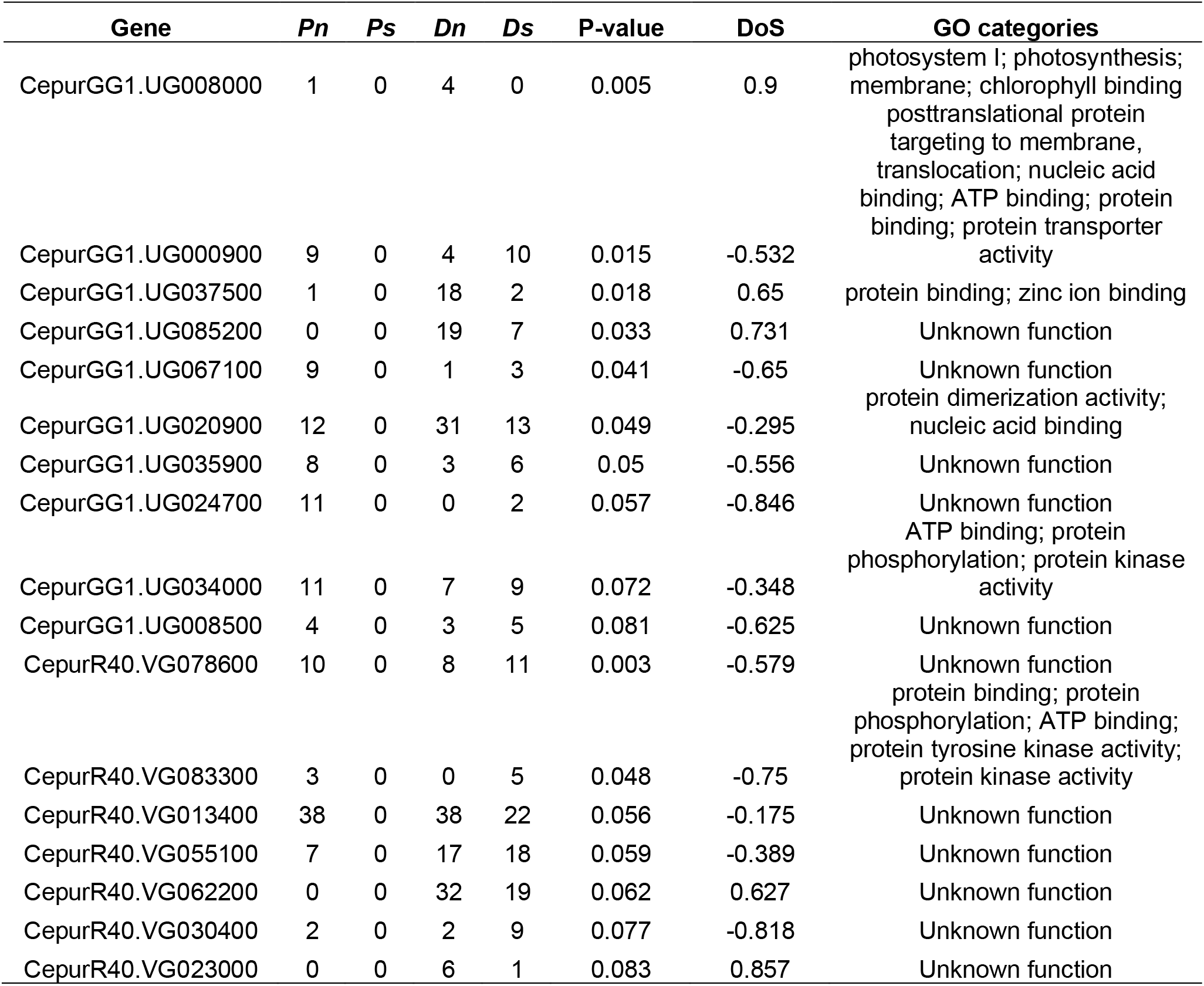
McDonald Krietman test results for sex-linked genes. Results shown here are significant in the MK test at p<0.1 (autosomal genes shown in Table S2). Non-synonymous polymorphism (*Pn*); Synonymous polymorphism (*Ps*); Non-synonymous divergence (*Dn*); Synonymous divergence (*Ds*); Direction of Selection (DoS); Gene Ontology (GO).

To test for differences in mutation rate, we calculated *dS* and *dN* on one-to-one U-V orthologs. We found both *dS* and *dN* were higher for V-linked genes (Mann Whitney U, *dN* p=0.044; *dS* p=0.005; Figure 4B-C; Table S4).

### Comparing empirical data to simulations

To determine if population density and sex-ratio bias could explain the observed patterns of *θ* we found in *C. purpureus*, we first calculated the variance in reproductive success in males (*V_m_*) and females (*V_f_*) that would be necessary to explain our empirical results and compared these values to those seen in our simulations. Specifically, for equal sex ratios we compared the ratios of *V_m_/V_f_* (*α*) and *V_m_/V_f_+V_m_* (*β*) described above. From our empirical data, we found that *α* and *β* were very similar values, with *α* = 1.36 (*α*= 1.62 if *μ_V_* was 1.2 times greater than *μ_U_* based on *dS*) and *β* = 1.0. In contrast, for all simulated densities, *α* was much larger than *β* (Figure 5A). Furthermore, our empirically observed *α* values were only seen in our simulations with low densities (~0.09–0.125), but the highest-density simulations were the ones that best approximated the empirically calculated *β* values.

**Figure 5.**
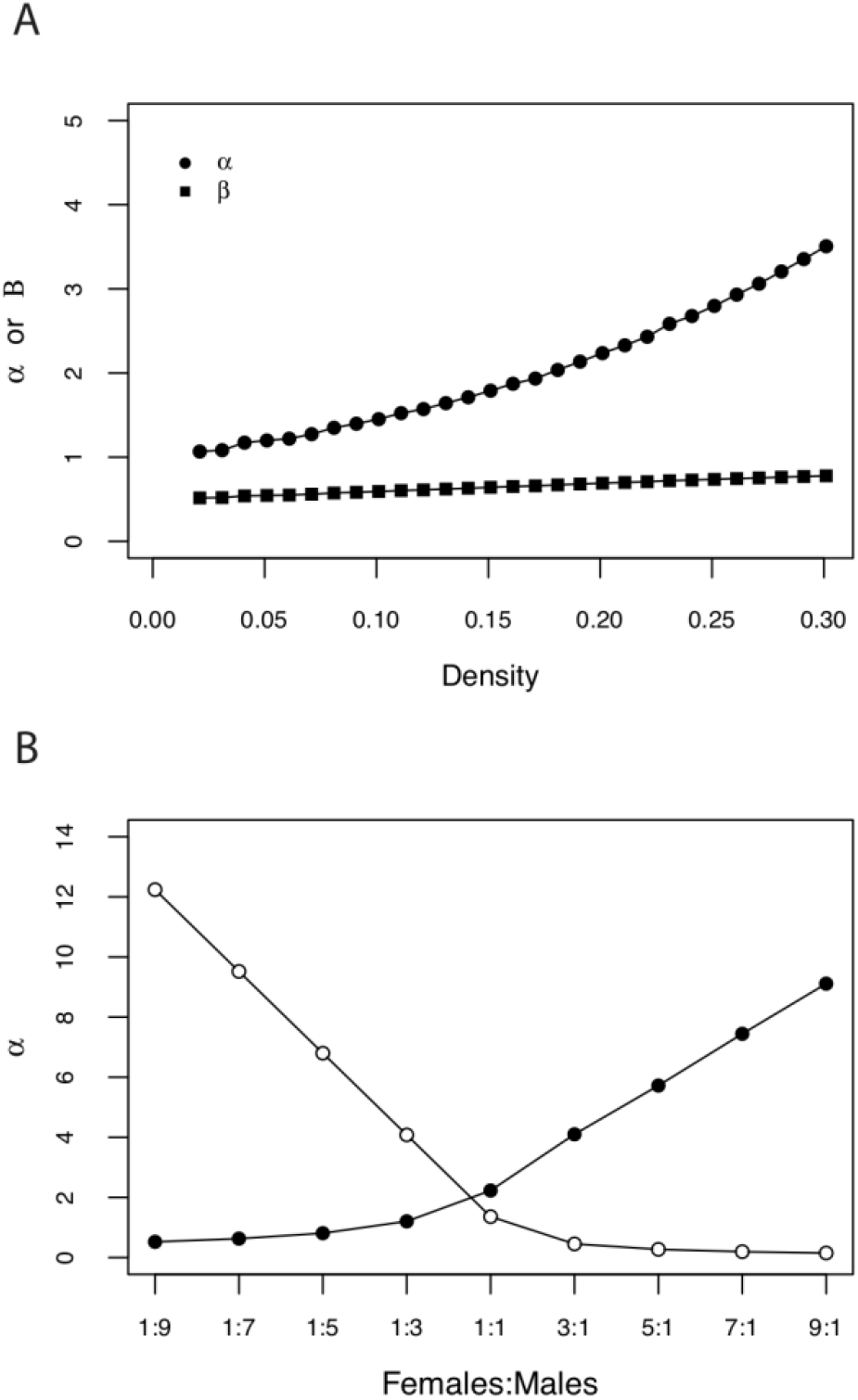
A) Simulation results for the effect of density on the two ratios discussed in the main text. The ratio between variances in reproductive success of males and females (α) and the variance in reproductive success of males divided by the variance in reproductive success of males plus the variance in reproductive success of females (β). The patterns plotted here hold for different populations and arena sizes. Each point is the mean value given by 100 runs of the simulation. Empirically calculated values were *α*= 1.36 and *β* = 1.0. B) The effect of sex ratio (females:males) on the ratio between variances in reproductive success of males and females (α). Filled black points show results of simulations (with a density of 0.2) while white points show the empirically calculated values. Each black point is the mean value given by 100 runs of the simulation.

For unequal sex ratios, the assumptions necessary to derive the equation for *β* do not hold, so we instead compared the ratios of *α* and *γ* described above. The combinations of sex ratio and density that best explained the observed values of *α* poorly predicted the observed *γ* values. For instance, at density of 0.2, our simulations best explained the observed *α* value when there was a slightly male-biased sex ratio (Figure 5B). However, in such scenarios, the simulated *γ* value was much larger than the *γ* values calculated from our data (e.g., for a simulated sex ratio of three males to each female, *γ* = 5.6x 1^−4^ while the value calculated for our data is *γ* = 8.3310^−9^). None of our simulations observed a *γ* value low enough to explain our data, but the closest values occurred at highly male-biased sex ratios.

## Discussion

Fertilization in many eukaryotes is achieved through the union of a small, motile sperm, and a large, retained egg. While the individuals that bear these gametes may experience different patterns of selection or embark on different patterns of migration, the consequences of life history differences between sexes alone may be sufficient to generate nucleotide diversity differences between sperm-producing males and egg-producing females. Here we used simulations to generate demographically-informed expectations for patterns of polymorphism in an idealized sperm-casting species and compared those expectations to estimates from whole-genome resequence data in the moss *C. purpureus*. We found the demographically-naive expectations for U and V chromosome-to-autosome ratios of nucleotide diversity based on the infinite-sites model were only accurate at the very lowest simulated population densities, and they failed to account for levels of sex-ratio bias observed in natural populations. We also found that our empirical estimates of sex chromosome and autosome nucleotide diversity could be explained by neither mutation rate variation nor other demographic processes, suggesting that selection must contribute to shaping variation on the *C. purpureus* sex chromosomes (Ellegren, 2009; Sayres, 2018).

### Sex-biased *N_e_* in anisogamous sperm-casting species

Estimates of *N_e_* can differ between females and males for numerous reasons, most dramatically as a consequence of processes that increase the variance in male reproductive success (i.e., female mate choice, male-male competition), but also because life-history differences may influence sex-specific patterns of migration or age structure within populations. Here we use simulations to show that in sperm-casting species, like some sessile marine animals and many plants, heterogeneity in the spatial distribution of females and males can generate a dramatic increase in the variance in male reproductive success. We show that this effect is strongly dependent upon the density of individuals. At low densities, the estimates of *N_e_* were relatively close to the estimates based on the infinite-sites model. The number of females and males that were near enough to mate was relatively low, and similar proportions of females and males contributed to the next generation. As density increased, more males mated multiple times, increasing the variance in male reproductive success and decreasing the male *N_eV_*. At high densities, the female *N_eU_* approached the census size, far exceeding both *N_eV_* and even *N_eA_*. This seemingly counterintuitive result stems from the fact that nearly all females reproduced, but many males contributed to multiple matings. The *N_eA_* is constrained to be between the male and female values, because half of the autosomes are inherited from each sex.

It is important to note that our simulation results relied on specific assumptions about the direct links among sex chromosomes, anisogamy, and life history. For example, we assumed that all individuals with a U-chromosome produced eggs and all individuals with a V-chromosome produced sperm, although it is well-known that these assumptions are violated in many systems (Ming, Bendahmane, & Renner, 2011). In *C. purpureus* many individuals do not produce gametangia under permissive laboratory conditions (J. Shaw & Beer, 1999). We also assumed that the egg-producing sex (females) and sperm-producing sex (males) each have specific, invariant life histories: females can only mate once and males can mate many times (eight is the maximum number of mating events possible in our simulations). While these provide a reasonable approximation of the life history of *C. purpureus* and other sperm-casting species, we caution that these results cannot be applied uncritically to other anisogamous species (see (Sarah Blaffer Hrdy, 1986; S. B. Hrdy, 1981; Tang-Martínez, 2016)).

How important the density-dependent effects on male variance in reproductive success are in sperm-casting species in nature depends upon how many distinct genotypes lie within the radius of sperm dispersal. Many bryophytes exhibit largely clonal growth, meaning the effective density of genotypes could be quite low, and the infinite-sites estimate may be a reasonable approximation (Bisang & Hedenäs, 2005; Clarke, Ayre, & Robinson, 2009). However, despite their capacity for clonal growth, small samples (<1cm) of *C. purpureus* can contain numerous distinct genotypes (McDaniel & Shaw, 2005). Given that fertilization distances in mosses exceed this measure (Jonathan Shaw & Goffinet, 2000; Longton & Re, 1976) particularly if transported by microarthopods (Cronberg et al., 2006; Rosenstiel et al., 2012; Shortlidge et al.,2020), the effective density may be quite high (i.e., the number of females that a given male can mate with exceeds the limit of eight imposed in our simulation). Thus, the infinite-sites expectations for *N_e_* may be quite far off. The mismatch between infinite-sites model and reality may be worse with female-biased sex ratios, which is common in bryophytes (Bisang & Hedenäs, 2005).

The decrease in *N_e_* for sperm-transmitted genes, relative to the *N_e_* for egg-transmitted genes, may profoundly decrease the strength of natural selection relative to genetic drift for male traits (Charlesworth, 2009). The decrease in efficacy of natural selection will likely be the most acute on a male V chromosome, where transmission is exclusively through sperm. However, the effects of anisogamy may influence the strength of selection on autosomal variants that have different fitness effects on females and males (i.e., sexually antagonistic alleles), effectively tipping the scales in favor of female-beneficial autosomal alleles. This process could even act on hermaphroditic sperm-casting species (Abbott, 2011), like many mosses, by weakening selection on allelic variants that promote male functions, such as sperm production, relative to selection on female functions related to egg production. Our haploid model does not allow us to make quantitative predictions about diploid systems, but male traits in seed plants may experience lower *N_e_* if some pollen donors fertilize multiple seeds in a population.

### Linking demographic models and mutation rate to patterns of nucleotide diversity

It is widely known that various demographic processes can generate variation in nucleotide diversity between autosomes and sex chromosomes, in particular processes that increase male variance in reproductive success (Charlesworth, 2009). Here we introduce three ratios of the sex-specific variance *N_e_*, *α* (variance in male reproductive success:female reproductive success), *β* (variance in male reproductive success:variance female reproductive success plus variance in male reproductive success), and *γ* (same as *β*, but the variances in the denominator are scaled by sex-specific population size), calculated from our simulations. These ratios can be expressed in terms of the quantity *θ*, which we estimated from the DNA sequence data from isolates of *C. purpureus*. The values of *α* and *β* generated from the simulations are clearly inconsistent with the nucleotide diversity patterns in *C. purpureus*. First, the *β*= 1 value that we calculated from the resequence data requires no variance in female reproductive success, which is inconsistent with mesocosm experiments in *C. purpureus* (Shortlidge et al., 2020) and field-collected data in *Sphagnum* (Johnson & Shaw, 2016). Similar *β* values were only observed at the highest population densities in our simulations. In contrast, the *α* values we estimated from the polymorphism data were only observed in simulations with low population densities. No simulated densities with an equal sex ratio produced values of both *α* and *β* near to those calculated from the nucleotide diversity patterns in *C. purpureus*.

Similarly, incorporating sex-ratio variation into our simulations failed to produce a better fit to the nucleotide diversity data. The closest *γ* values occurred at highly male-biased sex ratios, which are almost never observed in mosses (Baughman et al., 2017; Bisang et al., 2019; Bisang & Hedenäs, 2005) and in *C. purpureus* in particular (Eppley et al., 2018; Norrell et al.,2014; A. J. Shaw & Gaughan, 1993). In essence, the values for *θ_U_* and *θ_V_* are too low, relative to the autosomal *θ*, to conform to the demography alone model. In particular, the values for *θ_U_* were expected to equal or exceed the autosomal values under reasonable population density parameters, but instead the empirical *θ_U_* values were nearly as low as the *θ_V_* values. We return to this observation below.

We also found no evidence that the differences in *θ* between sex chromosomes was the result of an elevated mutation rate in females. In fact, both *dS* and *dN* were higher on V-linked genes (Mann Whitney U, *dN* p=0.044; *dS* p=0.005; Figure 4B-C), the opposite of the pattern that would explain the higher *θ* on the U chromosome compared to the V. We did recover the expected lower nucleotide diversity on the chloroplast (Table 1), consistent with other plants (D.R. Smith, 2015; Wolfe et al., 1987) suggesting that the elevated male mutation rate is unlikely to be an artifact of our sampling scheme.

Male-biased mutation rates are widely observed in animals and some seed plants (Ellegren & Fridolfsson, 1997; Whittle & Johnston, 2002; Wilson Sayres & Makova, 2011). Mutations presumably arise as DNA replication errors, and sperm production requires many more cell divisions than egg production (W. H. Li, Yi, & Makova, 2002). Because the germ line is sequestered in animals, somatic mutations do not contribute to differences between the sexes. Instead, only differences in cell divisions to produce gametes differ between the sexes. In contrast, plants have an open developmental program, in which many rounds of cell division precede the formation of gametes, potentially moderating the contribution of anisogamy to mutation-rate variation. However, males in mosses certainly make thousands of sperm in each antheridium (Garbary, Renzaglia, & Duckett, 1993). In the hermaphroditic moss, *Physcomitrium patens*, one antheridium (of ~10) made between 150 and 200 sperm cells (Horst & Reski, 2017), but this species is likely on the lower end of the distribution (Garbary et al., 1993). Thus, we conservatively estimate a *C. purpureus* male undergoes seven to 15 more rounds of cell division during sperm production than a female experiences in egg production, potentially enough to increase the male-mutation rate. Resequence data from known pedigrees is one way to independently evaluate the difference between male and female mutation rates, but such data are currently unavailable.

### Selection lowers nucleotide diversity on U and V sex chromosomes

The relationship we observed between autosomal and V-linked nucleotide diversity plausibly could reflect elevated variance in male reproductive success, consistent with field studies and experimental mesocosms (Johnson & Shaw, 2016; Shortlidge et al., 2020). However, none of the simulations using biologically reasonable conditions explained the low U-linked nucleotide diversity (U/A=~1/3). It is therefore likely that the low nucleotide diversity on at least the U is caused by recent linked selection, which is widely expected to be common in non-recombining regions, like sex chromosomes (J. M. Smith & Haigh, 1974). Indeed, Tajima’s D values on both the U and the V were uniformly negative and lower than autosomal values (Figure 3; Table 1), suggesting the patterns of nucleotide variation do not reflect neutral-equilibrium processes. The *C. purpureus* UV sex chromosomes are large (>100 Mb each), completely non-recombining, and gene rich (>3,400 genes annotated to both (S. B. Carey et al., 2020)) meaning these regions provide large targets for the evolution of beneficial or deleterious mutations, increasing the probability of selective sweeps and background selection (Bachtrog, 2008).

Identifying the relative importance of selective sweeps and background selection in reducing nucleotide diversity remains an important challenge. The *C. purpureus* sex chromosomes demonstrably experience weaker purifying selection than autosomes, based on measures of codon bias (effective number of codons, frequency of optimal codon, and GC content of the third synonymous position) and in protein evolution (*dN/dS*) (S. B. Carey et al.,2020). Carey et al. (2020) found that, of the 330 U and V-linked genes examined, ~25% had lower *dN/dS* than the autosomal average (~0.14), suggesting these genes still experience strong selective pressure against deleterious mutations (Chibalina & Filatov, 2011). However, codon bias and *dN/dS* were indistinguishable between the sexes, suggesting that different levels of purifying selection are unlikely to explain the decreased V/U ratio we found. Moreover, while the U and V-linked genes with higher *dN/dS* values suggest faster rates of protein evolution than the autosomes, using this approach it is unclear whether the genes are evolving faster due to the relaxation of purifying selection or by positive selection.

Because our sampling spanned the globe, we used the distantly-related Chilean isolates as an outgroup for the Northern Hemisphere populations in divergence-polymorphism tests of selection. Using the MK test, we also found evidence of non-neutral evolution in several sex-linked genes (Figure 4A; Table 2). From the DoS test we found the U has genes that are experiencing positive selection, while others have relaxed purifying selection (at p<0.05; Figure 4A). On the V we found genes with evidence of relaxed purifying selection (at p<0.05), though two genes were marginally significant that suggest positive selection (p≃0.06 and 0.08). Some of the genes that showed evidence of selection are involved in cellular transport (Table 2). In mosses, the sporophyte (i.e., the diploid embryo) is nutritionally dependent on the maternal plant throughout its entire lifespan (Ligrone, Duckett, & Renzaglia, 1993), which is costly to the maternal gametophyte (Ehrlén, Bisang, & Hedenäs, 2000; Stark, Brinda, & McLetchie, 2009). The male with which a female mates has a significant effect on sporophyte development including sporophyte height, spore number, and quality of spores (Shortlidge et al., 2020), suggesting paternal genotype can drive these differences. However, a sporophyte from a male with a more extractive genotype may instead be selectively aborted by the female in preference for another, less extractive offspring. In fact, female mosses have been shown to abort their offspring if conditions are unfavorable (Stark, 2002; Stark, Mishler, & McLetchie, 2000; Stark & Stephenson, 1983). These forms of sexual conflict can generate the signatures of selection detectable by the MK test, and could reduce nucleotide diversity on both the U and V chromosomes, consistent with what we find in *C. purpureus*. Though we should point out that the polymorphism-based tests are underpowered to detect deviations from neutrality in regions of low *N_e_* (like non-recombining sex chromosomes) and limited divergence between our outgroup (Parsch, Zhang, & Baines, 2009).

The discrepancy between the infinite-sites expectations and the measured nucleotide diversity that we found in *C. purpureus* sex chromosomes, in particular the U, are qualitatively different from those in other plant systems, in spite of the fact that all possess multicellular, gametophytes with haploid gene expression. *In Rumex hastatulus*, nucleotide diversity of X-linked genes was ~⅘ of autosomal diversity, higher than neutral equilibrium expectations, potentially because of the female-biased sex ratios in the species (Hough, Wang, Barrett, & Wright, 2017). In contrast, the Y-linked genes were ~1/50 that of autosomal diversity, a result attributed to purifying selection (Hough et al., 2017). In *Silene latifolia*, Y-linked genes had ~1/20 the nucleotide diversity of autosomes, whereas X-linked genes were close to equilibrium expectations (~3/4), also suggesting the role of selection on the Y (Qiu, Bergero, Forrest, Kaiser, & Charlesworth, 2010). The X chromosome in papaya was found to have lower than expected nucleotide diversity, likely driven by a selective sweep, like what we found on the U in *C. purpureus* (VanBuren et al., 2016). In the brown algae *Ectocarpus*, the only other UV system to look at nucleotide diversity to date, the sex chromosomes have ~1/2 *N_e_* of autosomes, consistent with infinite-sites expectations for neutral-equilibrium conditions (Avia et al., 2018).

Together these results support the long-standing notion that positive selection can dramatically decrease nucleotide diversity in non-recombining regions (Begun & Aquadro, 1992; Charlesworth & Charlesworth, 2000; Lercher & Hurst, 2002). The dramatic difference between the simulated and empirical estimates for *N_e_* for the *C. purpureus* U provide a very clear illustration of this effect. Our simulations also suggest that density, mating system, and factors that influence the variance in male reproductive success may also be confounded with selection in analyses of Y chromosome polymorphism.

The geographic distribution of U and V-linked variants may provide insight into other forms of selection shaping sex chromosomea riation. Of course, differential migration between the sexes can affect *N_e_* (Ellegren, 2009; Goudet, Perrin, & Waser, 2002; Webster & Wilson Sayres, 2016). However, at a regional scale among several eastern North American populations, and with a smaller data set, population structure measured by *F*_ST_ (Sewall Wright, 1949) was equivalent between the sexes in *C. purpureus* (McDaniel, Neubig, et al., 2013). Importantly, the U and V *F*_ST_ among these populations was lower than the autosomal *F*_ST_. This pattern suggests that sex chromosome variants are fit across the region, while autosomal alleles may be more likely to experience local adaptation, and therefore show elevated *F*_ST_ values. Here, we found that *F*_ST_ on the sex chromosomes on the continental scale between eastern and western North American populations were also equivalent between the sexes, but exceeded the autosomal values (*F*_ST_ Autosomes=0.202, U=0.372, V=0.375; Table 1). At the continental scale, autosomal alleles are exchanged among these populations while migrant sex chromosomes are not. Patterns of interfertility (McDaniel, Willis, & Shaw, 2008) and preliminary gene tree analyses suggest that the eastern and western North American populations may represent partially reproductively isolated species. The low *F*_ST_ at the regional scale coupled with higher *F*_ST_ at the continental scale suggests that sex chromosome differentiation may occur at the scale of species boundaries, rather than the scale of local adaptation. This inference is consistent with data from *Drosophila* and primates showing that more accurate phylogenies are inferred from non-recombining regions, like sex chromosomes (Pease & Hahn, 2013).

Together, these results highlight the utility of UV sex chromosomes as models for understanding the roles of sex-specific evolutionary processes in genome evolution. The demographic model that we present shows that small male *N_e_* may be a critical challenge facing dioecious species, a potentially important factor to explain features of mating-system variation in bryophytes, including frequent transitions from dioecy to hermaphroditism and the evolution of dwarf males (Hedenäs & Bisang, 2011; McDaniel, Atwood, & Burleigh, 2013). Similar to other eukaryotic lineages, the increase in variance in reproductive success is also correlated with a modestly increased mutation rate. Perhaps the most striking result is the decrease in U/A ratio of *N_e_*, relative to the expectations based on simulations. The difference between the simulated and empirical values strongly suggests that the U sex chromosome experiences frequent selective sweeps, an inference with some independent support from frequency spectrum and codon-based molecular evolutionary analyses. In sum, these data highlight the challenges of conducting analyses of single evolutionary forces, in isolation, without considering their joint effects.

## Acknowledgements

We thank the U.S. Department of Energy Joint Genome Institute for pre-publication access to the *Ceratodon purpureus* genome used in this study and Leslie Kollar for helpful feedback on this manuscript. The University of Florida (UF) HiPerGator provided vital technical support throughout the project. This work was supported by NSF DEB-1541005 and start-up funds from UF to SFM. The work conducted by the U.S. Department of Energy Joint Genome Institute was supported by the Office of Science of the U.S. Department of Energy under Contract No. DE-AC02-05CH11231.

## Data Accessibility

Requests for the *C. purpureus* lines in this manuscript should be addressed to. The *C. purpureus* Illumina resequencing data can be found under NCBI BioProjects listed in Table S1. The *C. purpureus* chloroplast assembly can be found on NCBI under (in progress). Code for the life history simulations can be found at https://github.com/JimmyPeniston/Moss_NE_simulations and the population genetic analyses can be found at https://github.com/sarahcarey/Ceratodon_popgen.

## Author Contributions

SBC, JHP, and SFM designed the research; ACP, AL, DB, KL, CD, KB, JG, JS, and SFM performed the molecular biology and sequencing; SBC, MK, JJ, and JS performed the bioinformatics; JHP developed the model; SBC performed the population genetic analyses; SBC and JHP created the visualizations; SBC, JHP, and SFM wrote the original draft and edited the manuscript.

