## Supplementary Materials for "Novel insights into joint estimations of demography, mutation rate, and selection using UV sex chromosomes"

**This PDF file includes:**

Appendix 1 Parts 1-2

Supplemental Tables 1-4

Supplemental Figure 1

**Appendix 1**

**Part 1: Inbreeding effective population sizes**

The inbreeding effective population size is the number of individuals that an idealized population would need in order for the probability of identity by descent of two randomly chosen alleles to match that of the census population (Crow & Kimura, 1970). Specifically, if *Pt* is the probability that two randomly chosen gametes are descended from the same individual in previous generation, the inbreeding effective population size is

$$N_e = \frac{1}{P_t}$$

Here we will derive the inbreeding effective population size (hereafter "effective population size" or $N_e$) for different chromosomes (sex chromosomes *versus* autosomes) in a haploid-dioecious system. In particular, we will focus on how variation in reproductive success and sex ratio affects effective population size. While many other researchers have used the same or similar equations, the derivation is usually only shown for diploid populations. We closely follow the derivation presented by Crow and Kimura (1970) but make the appropriate changes to fit our life history assumptions.

*Effective population size of U and V chromosomes*

In a haploid-dioecious system, the sex chromosomes (U and V) are sex specific. Therefore, the effective population sizes of the U and V chromosomes can be modeled as independent haploid populations.

Let $N_t$ be the number of haploid adults in the population at time *t* (the census population size) and let $k_i$ be the number of successful gametes from the $i^{th}$ parent in generation $t-1$. Therefore, the total number of gametes in generation *t* is $N_{t-1}\bar{k}$, where $\bar{k}$ is the mean number of gametes an individual contributes to the next generation. The number of ways to randomly select two gametes from the $N_{t-1}\bar{k}$ gametes in the population is given by

$$\binom{N_{t-1}\bar{k}}{2},$$

and the number of ways in which two gametes came from the same parent is

$$\sum_i \binom{k_i}{2}.$$

Therefore, the probability that two randomly selected gametes in generation *t* are descended from the individual is

$$P_t = \frac{\sum_i \binom{k_i}{2}}{\binom{N_{t-1}\bar{k}}{2}} = \frac{\sum_i k_i(k_i - 1)}{N_{t-1}\bar{k}(N_{t-1}\bar{k} - 1)}.$$

The variance in the number of gametes contributed per individual (variance in reproductive success) is defined as

$$V_k = \frac{\sum_i (k_i - \bar{k})^2}{N_{t-1}} = \frac{\sum_i k_i^2}{N_{t-1}} - \bar{k}^2.$$

If we note that the above equation can be rearranged in the form

$$\sum_i k_i^2 = N_{t-1}(V_k + \bar{k}^2),$$

and that $N_{t-1}\bar{k} = \sum_i k_i$, equation A4 can be rewritten as

$$P_t = \frac{\left[N_{t-1}(V_k + \bar{k})\right] - (N_{t-1}\bar{k})}{N_{t-1}\bar{k}(N_{t-1}\bar{k} - 1)}.$$

Noting that in a haploid system $N_{t-1}\bar{k} = N_t$, this can be simplified to

$$P_t = \frac{N_{t-1}(V_k + \bar{k}^2 - \bar{k})}{N_t(N_{t-1}\bar{k} - 1)}$$

$$P_t = \frac{N_{t-1}(V_k + \bar{k}^2 - \bar{k})}{N_t\bar{k} - (N_t / N_{t-1})}$$

$$P_t = \frac{N_{t-1}(V_k + \bar{k}^2 - \bar{k})}{N_t\bar{k} - (N_t\bar{k} / N_t)}$$

$$P_t = \frac{V_k + \bar{k}(\bar{k} - 1)}{\bar{k}(N_t - 1)}.$$

Thus, the inbreeding effective population size ($1/P_t$) is

$$N_e = \frac{\bar{k}(N_t - 1)}{V_k + \bar{k}^2(\bar{k} - 1)}.$$

If we assume that the population size remains constant, $\bar{k} = 1$, and thus the effective population size of the U and V chromosomes is given by

$$N_e = \frac{N_t - 1}{V_K}.$$

*Effective population size of autosomes*

In haploid-dioecious systems, there is one copy of each autosome in every individual. Despite adults being haploid, each individual is still created during a fertilization event that involves the fusion of egg and sperm (this diploid phase then produces haploid spores which form into adults; (Bachtrog et al., 2011)). We refer to the egg donors as females and the sperm donors as males and assume that all individuals are either male or female. Therefore, in the absence of selection, both sexes equally contribute to the next generation. The probability that two randomly selected autosomes in the population are derived from the same female parent is

$1/4N_{ef}$ and the probability that they were selected from the same male $1/4N_{em}$, where $N_{ef}$ is the effective population size of females and $N_{em}$ is the effective population size of males. Therefore, the probability the probability that two randomly selected autosomes in the population were derived from the same individual is

$$\frac{1}{4N_{ef}} + \frac{1}{4N_{em}} = \frac{1}{N_{eA}},$$

which can be rewritten in the common form (Lande & Barrowclough, 1987)

$$N_{eA} = \frac{4N_{ef}N_{em}}{N_{ef} + N_{em}},$$

where $N_{eA}$ is the effective population size of the autosomes. In order to directly relate the autosomal effective population size, to that of the sex chromosomes, we make the simplifying assumption that all females (egg donors) carry a U chromosome and all males (sperm donors) carry a V chromosome. Thus,

$$N_{eA} = \frac{4N_{eU}N_{eV}}{N_{eU} + N_{eV}},$$

where $N_{eU}$ and $N_{eU}$ are given by equation A9.

**Part 2: Number of segregating sites**

Here we will derive the number of neutral alleles maintained in a finite population. Our definition is based on the infinitely many alleles model, which assumes that every mutation leads to a new allele of an entirely novel type (Ewens, 2016; Malécot, 1948). We follow the derivation of Crow and Kimura (1964, 1970), but make the appropriate changes to fit a haploid system (Crow & Kimura, 1970; Kimura & Crow, 1964).

In a population of effective size, $N_e$, the inbreeding coefficient is given by

$$f_t = \frac{1}{N_e} + \left(1 - \frac{1}{N_e}\right) f_{t-1}.$$

The first term is the probability that the alleles are identical because they are derived from the same individual and the second term of alleles from two different individuals in the previous generation being identical. If we let $\mu$ be the mutation rate, the probability that two identical alleles will remain so the next generation is given by $(1-\mu)^2$, which is the probability that neither mutated. Therefore,

$$f_t = \left[\frac{1}{N_e} + \left(1 - \frac{1}{N_e}\right) f_{t-1}\right](1-\mu)^2.$$

Assuming that an equilibrium is reached such that $f_t = f_{t-1}$,

$$f_t = \frac{(1-\mu)^2}{N_e - (N_e - 1)(1-\mu)^2}.$$

If we assume that $\mu$ is very small, we can ignore the terms containing $\mu^2$. Therefore,

$$f_t = \frac{(1-2\mu)}{2N_e\mu - 2\mu - 1} \approx \frac{1}{2N_e\mu - 1}.$$

The number of segregating alleles is given by $\theta = 1/f_t$ (Crow & Kimura, 1970), therefore,

$$\theta = 2N_e\mu - 1.$$

Following previous work (e.g. (Watterson, 1975)), we approximate this as

$$\theta = 2N_e\mu.$$

### Supplemental Tables

**Table S1. *Ceratodon purpureus* isolate information, library information, and mapping percentages.** For isolates that were sequenced across two lanes, there are two JGI sequence files, and the mapping values are shown in the same order as these files. The first files were sequenced using version 3 chemistry on a HiSeq2000. The second files (if present) were sequenced using version 4 chemistry on a HiSeq2500.

| Isolate | Location | Library | JGI sequence file(s) | Total reads | Sex | BWA reads mapped | BWA % mapped | NGM reads mapped | NGM % mapped | Coverage | NCBI SRA | NCBI BioProject |
|---|---|---|---|---|---|---|---|---|---|---|---|---|
| Alaska 21.1.1 M1 | Yukon, Alaska, USA | WPUH | 8032.4.89493.ACAGTG.anqdp.fastq.gz; 8372.1.99482.ACAGTG.anqdp.fastq.gz | 53566984; 89190836 | Male | 41414343; 70257445 | 77.31; 78.77 | 41499167; 70260545 | 77.47; 78.78 | 36.35888505 | SRP083255 | PRJNA258679 |
| Alaska 21.1.8 F1 | Yukon, Alaska, USA | WPUG | 8032.3.89495.TGACCA.anqdp.fastq.gz | 71566146 | Female | 50128990 | 70.05 | 50135077 | 70.05 | 17.66073196 | SRP083242 | PRJNA258678 |
| Ant 1 | Casey Base, Antarctica | WPWS | 8066.4.90145.ACTGAT.anqdp.fastq.gz; 8372.5.99498.ACTGAT.anqdp.fastq.gz | 71831400; 104143574 | Female | 66929671; 96909345 | 93.18; 93.05 | 66770712; 96931815 | 92.95; 93.08 | 54.51409029 | SRP083369 | PRJNA258684 |
| Au 1 | Australia | WPWA | 8066.2.90149.GTAGAG.anq | 33380834; | Female | 31543453; 52308754 | 94.5; 94.57 | 31508238; | 94.39; 94.55 | 29.959877 | SRP083258 | PRJNA258682 |

| | | | | | | | | | | | | |
|---|---|---|---|---|---|---|---|---|---|---|---|---|
| | | | dp.fastq.gz; 8372.3.99490.GTAGAG.anqdp.fastq.gz | 55310268 | | | | 52297424 | | | | |
| Au 2 | Australia | WPWB | 8066.2.90149.GTCCGC.anqdp.fastq.gz; 8372.3.99490.GTCCGC.anqdp.fastq.gz | 56907848; 72107000 | Female | 53642093; 68117015 | 94.26; 94.47 | 53454828; 67977993 | 93.93; 94.27 | 35.86396963 | SRP083344 | PRJNA258683 |
| Chile 1.1 M1 | is Navarino, Magellanes, Chile | WPUB | 8032.3.89495.CGATGT.anqdp.fastq.gz | 68407526 | Male | 64900045 | 94.87 | 64754593 | 94.66 | 20.88551582 | SRP083246 | PRJNA258681 |
| Chile 2.12 F1 | is Navarino, Magellanes, Chile | WPUC | 8032.3.89495.TTAGGC.anqdp.fastq.gz | 53131298 | Female | 50359560 | 94.78 | 50210782 | 94.5 | 16.58103519 | SRP083244 | PRJNA258680 |
| Dur 13.8.1 M3 | Durham, NC, USA | WPWN | 8066.3.90147.CGTACG.anqdp.fastq.gz; 8372.4.99494.CGT | 62367540; 80373166 | Male | 26370976; 38802721 | 42.28; 48.28 | 27916469; 39819881 | 44.76; 49.54 | 22.39103426 | SRP083353 | PRJNA258700 |

| | | | | | | | | | | | | |
|---|---|---|---|---|---|---|---|---|---|---|---|---|
| | | | ACG.anqdp.fastq.gz | | | | | | | | | |
| Dur 13.8.10 F3 | Durham, NC, USA | WPWH | 8066.3.90147.GTTTCG.anqdp.fastq.gz; 8372.4.99494.GTTTCG.anqdp.fastq.gz | 22844040; 30719068 | Female | 10439930; 15374593 | 45.70; 50.05 | 10938383; 15702446 | 47.88; 51.12 | 9.267406989 | SRP083355 | PRJNA258999 |
| Dur 16.6.2 F1 | Durham, NC, USA | WPUU | 8032.6.89492.CTTGTA.anqdp.fastq.gz | 54274564 | Female | 30375562 | 55.97 | 31138343 | 57.37 | 11.46888793 | SRP083257 | PRJNA259000 |
| Dur 19.2.8 M1 | Durham, NC, USA | WPUT | 8032.5.89498.GGCTAC.anqdp.fastq.gz; 8372.2.99486.GGCTAC.anqdp.fastq.gz | 58936240; 79359484 | Male | 24865353; 34044029 | 42.19; 42.9 | 26035379; 35192938 | 44.18; 44.35 | 21.58729629 | SRP083277 | PRJNA259004 |
| Dur 4.13.5 M2 | Durham, NC, USA | WPWC | 8066.2.90149.GTGAAA.anqdp.fastq.gz; 8372.3.99490.GTGAAA.anqdp.fastq.gz | 30159950; 39344502 | Male | 9197069; 13488542 | 30.49; 34.28 | 10150271; 14238904 | 33.65; 36.19 | 8.250311023 | SRP083345 | PRJNA258998 |
| Dur 4.13.7 F2 | Durham, NC, USA | WPWG | 8066.3.90147.GTGGCC.anq | 36227946; | Female | 34793594; 63733601 | 96.04; 96.28 | 34695498; | 95.77; 96.08 | 33.41230438 | SRP083346 | PRJNA258997 |

| | | | | | | | | | | | | |
|---|---|---|---|---|---|---|---|---|---|---|---|---|
| | | | dp.fastq.gz; 8372.4.99494.GTGGCC.anqdp.fastq.gz | 66192736 | | | | 63595215 | | | | |
| Equ E13.E3.1 F1 | Otavalo, Ecuador | WPUO | 8032.4.89493.CAGATC.anqdp.fastq.gz; 8372.1.99482.CAGATC.anqdp.fastq.gz | 43037242; 58510508 | Both; likely contamination; treated as female in analyses | 40688457; 55519415 | 94.54; 94.89 | 40548650; 55401302 | 94.22 94.69 | 30.33503249 | SRP083250 | PRJNA259003 |
| Equ E13.E3.14 M1 | Otavalo, Ecuador | WPUN | 8032.4.89493.GCCAAT.anqdp.fastq.gz; 8372.1.99482.GCCAAT.anqdp.fastq.gz | 62792852; 84853180 | Male | 59752425; 80872527 | 95.16; 95.31 | 59619574; 80816325 | 94.95; 95.24 | 43.33864 | SRP083249 | PRJNA259005 |
| Port 11 F2 | Portland, OR, USA | WPWP | 8066.4.90145.GGTAGC.anqdp.fastq.gz; 8372.5.99498.GGTAGC.anqdp.fastq.gz | 40777074; 77597512 | Female | 39141976; 74684319 | 95.99; 96.25 | 39038663; 74578679 | 95.74; 96.11 | 39.1924447 | SRP083386 | PRJNA259002 |

| | | | | | | | | | | | | |
|---|---|---|---|---|---|---|---|---|---|---|---|---|
| Port 4 M2 | Portland, OR, USA | WPWO | 8066.4.90145.GAGTGG.anqdp.fastq.gz; 8372.5.99498.GAGTGG.anqdp.fastq.gz | 41340320; 85029124 | Male | 35906922; 75413835 | 86.86; 88.69 | 35693948; 75116937 | 86.34; 88.34 | 39.34455715 | SRP083373 | PRJNA259001 |
| Port 7 F1 | Portland, OR, USA | WPUP | 8032.5.89498.GATCAG.anqdp.fastq.gz; 8372.2.99486.GATCAG.anqdp.fastq.gz | 50654928; 64887838 | Both; likely contamination; treated as female in analyses | 46722917; 60158619 | 92.24; 92.71 | 46528206; 60059437 | 91.85; 92.56 | 37.9327394 | SRP083248 | PRJNA259008 |
| Port 8 M1 | Portland, OR, USA | WPUS | 8032.5.89498.TAGCTT.anqdp.fastq.gz; 8372.2.99486.TAGCTT.anqdp.fastq.gz | 67128916; 77610510 | Male | 63834706; 73830223 | 95.09; 95.13 | 63593920; 73680857 | 94.73; 94.94 | 51.13283505 | SRP083266 | PRJNA259007 |
| Stl F1 | St. Louis, MO, USA | WPUZ | 8066.1.90146.CCGTCC.anqdp.fastq.gz | 54450460 | Female | 52970944 | 97.28 | 52871406 | 97.1 | 19.2362897 | SRP083275 | PRJNA259006 |
| Stl M1 | St. Louis, MO, USA | WPUY | 8066.1.90146.ATGT | 119507960 | Female | 115383064 | 96.55 | 115140901 | 96.35 | 37.48427802 | SRP083260 | PRJNA259011 |

| | | | | | | | | | | | | |
|---|---|---|---|---|---|---|---|---|---|---|---|---|
| | | | CA.anqdp.fastq.gz | | | | | | | | | |
| Uconn 12.2.4 F2 | Storrs, CT, USA | WPUX | 8032.6.89492.AGTTCC.anqdp.fastq.gz | 63657776 | Male | 61270485 | 96.25 | 61065064 | 95.93 | 21.90912441 | SRP083273 | PRJNA259010 |
| Uconn 15.12.12 F1 | Storrs, CT, USA | WPUW | 8032.6.89492.AGTCAA.anqdp.fastq.gz | 54501120 | Female | 52299472 | 95.96 | 52131707 | 95.65 | 18.07693506 | SRP083276 | PRJNA259009 |

**Table S2. Population genetic analyses by chromosome using the NGM mapper.** Results using the BWA mapper are presented in Table 1. Segregating sites (*S*); Wu and Watterson's theta ($\theta$); Pi ($\pi$).

| Chromosome | Total sites | $S$ | $\theta$ | $\pi$ | Tajima's D | $F_{ST}$ |
|---|---|---|---|---|---|---|
| 1 | 29001003 | 808503 | 0.0085 | 0.0084 | -0.0728 | 0.185 |
| 2 | 26629683 | 972416 | 0.0111 | 0.01 | -0.4713 | 0.25 |
| 3 | 25160467 | 750643 | 0.0091 | 0.0091 | -0.0418 | 0.2262 |
| 4 | 22785024 | 728381 | 0.0097 | 0.0097 | -0.0814 | 0.2731 |
| 5 | 19969424 | 623677 | 0.0095 | 0.01 | 0.1832 | 0.212 |
| 6 | 18980603 | 596309 | 0.0095 | 0.0092 | -0.2075 | 0.1703 |
| 7 | 17972677 | 518903 | 0.0088 | 0.0083 | -0.2721 | 0.1763 |
| 8 | 17567963 | 541017 | 0.0093 | 0.0092 | -0.1182 | 0.2045 |
| 9 | 17527894 | 611202 | 0.0106 | 0.0107 | 0.0094 | 0.1808 |
| 10 | 17229405 | 502411 | 0.0088 | 0.0088 | -0.0678 | 0.1994 |
| 11 | 16661191 | 577824 | 0.0105 | 0.0105 | -0.05 | 0.2416 |
| 12 | 16459275 | 510877 | 0.0094 | 0.0094 | -0.0884 | 0.2416 |
| Chloroplast | 105555 | 262 | 0.0007 | 0.0008 | 0.1889 | 0.1884 |
| U | 112179120 | 803882 | 0.0028 | 0.0028 | -0.2979 | 0.3814 |
| V | 110524308 | 554087 | 0.0019 | 0.0018 | -0.6574 | 0.3894 |

**Table S3. McDonald Kreitman results for autosomal genes.** Results shown here are significant in the MK test at p<0.1 (autosomal genes shown in Table S2). Non-synonymous polymorphism (*Pn*); Synonymous polymorphism (*Ps*); Non-synonymous divergence (*Dn*); Synonymous divergence (*Ds*); Direction of Selection (DoS).

| Gene | *Pn* | *Ps* | *Dn* | *Ds* | P-value | DoS |
|---|---|---|---|---|---|---|
| CepurR40.7G057700 | 2 | 0 | 9 | 3 | 0 | 0.655 |
| CepurR40.2G244900 | 43 | 8 | 13 | 3 | 0 | 0.494 |
| CepurR40.3G233700 | 106 | 7 | 5 | 12 | 0 | -0.463 |
| CepurR40.1G042400 | 2 | 0 | 5 | 0 | 0 | 0.909 |
| CepurR40.1G195900 | 0 | 0 | 3 | 0 | 0 | 1 |
| CepurR40.1G021000 | 24 | 1 | 8 | 12 | 0 | -0.489 |
| CepurR40.2G052300 | 68 | 0 | 7 | 19 | 0.001 | -0.378 |
| CepurR40.10G017000 | 4 | 0 | 6 | 1 | 0.001 | 0.697 |
| CepurR40.6G047800 | 20 | 2 | 8 | 0 | 0.001 | 0.592 |
| CepurR40.9G097900 | 95 | 0 | 1 | 6 | 0.001 | -0.649 |
| CepurR40.7G096700 | 33 | 1 | 39 | 7 | 0.002 | 0.307 |
| CepurR40.4G204500 | 20 | 2 | 6 | 5 | 0.002 | -0.455 |
| CepurR40.2G254600 | 27 | 4 | 0 | 9 | 0.002 | -0.563 |
| CepurR40.2G237600 | 280 | 37 | 5 | 10 | 0.002 | -0.39 |
| CepurR40.3G006900 | 36 | 0 | 6 | 6 | 0.003 | -0.423 |
| CepurR40.1G157400 | 5 | 0 | 0 | 8 | 0.003 | -1 |
| CepurR40.8G072000 | 7 | 15 | 10 | 0 | 0.003 | 0.5 |
| CepurR40.8G184200 | 23 | 0 | 0 | 5 | 0.003 | -0.742 |
| CepurR40.1G241900 | 5 | 1 | 0 | 16 | 0.004 | -0.455 |
| CepurR40.2G082300 | 64 | 2 | 9 | 2 | 0.004 | 0.457 |
| CepurR40.3G011800 | 11 | 0 | 3 | 5 | 0.005 | -0.625 |
| CepurR40.1G012500 | 2 | 1 | 0 | 14 | 0.005 | -0.667 |
| CepurR40.3G167800 | 2 | 0 | 4 | 0 | 0.005 | 0.818 |
| CepurR40.3G011400 | 11 | 0 | 5 | 9 | 0.005 | -0.56 |
| CepurR40.5G111500 | 31 | 4 | 5 | 13 | 0.006 | -0.355 |
| CepurR40.3G247200 | 3 | 1 | 7 | 7 | 0.007 | 0.412 |
| CepurR40.1G054300 | 13 | 2 | 0 | 9 | 0.007 | -0.464 |
| CepurR40.5G142600 | 486 | 12 | 21 | 2 | 0.007 | 0.259 |
| CepurR40.5G121500 | 1 | 0 | 5 | 1 | 0.008 | 0.722 |
| CepurR40.1G077900 | 1 | 3 | 5 | 0 | 0.008 | 0.5 |
| CepurR40.10G029400 | 48 | 2 | 3 | 6 | 0.008 | -0.454 |
| CepurR40.11G015100 | 15 | 2 | 0 | 4 | 0.008 | -0.833 |
| CepurR40.9G152000 | 34 | 1 | 0 | 13 | 0.009 | -0.34 |
| CepurR40.9G145400 | 26 | 1 | 5 | 0 | 0.009 | 0.612 |
| CepurR40.7G104200 | 54 | 6 | 0 | 12 | 0.009 | -0.353 |
| CepurR40.8G072100 | 23 | 5 | 6 | 0 | 0.009 | 0.566 |

| | | | | | | |
|---|---|---|---|---|---|---|
| CepurR40.6G170700 | 6 | 0 | 2 | 6 | 0.01 | -0.75 |
| CepurR40.6G047000 | 14 | 2 | 1 | 5 | 0.01 | -0.611 |
| CepurR40.11G108100 | 39 | 9 | 0 | 6 | 0.01 | -0.534 |
| CepurR40.11G090300 | 60 | 0 | 6 | 9 | 0.01 | -0.369 |
| CepurR40.3G109300 | 21 | 2 | 12 | 1 | 0.01 | 0.411 |
| CepurR40.6G048800 | 19 | 4 | 5 | 0 | 0.01 | 0.642 |
| CepurR40.2G175200 | 32 | 1 | 0 | 4 | 0.01 | -0.711 |
| CepurR40.8G134900 | 8 | 4 | 6 | 16 | 0.011 | -0.394 |
| CepurR40.2G121000 | 63 | 2 | 1 | 3 | 0.011 | -0.625 |
| CepurR40.8G117000 | 16 | 0 | 1 | 6 | 0.011 | -0.584 |
| CepurR40.2G133800 | 5 | 1 | 5 | 0 | 0.011 | 0.688 |
| CepurR40.4G182600 | 32 | 3 | 0 | 3 | 0.011 | -0.8 |
| CepurR40.10G142100 | 12 | 0 | 8 | 0 | 0.011 | 0.5 |
| CepurR40.2G066000 | 16 | 1 | 8 | 1 | 0.011 | 0.479 |
| CepurR40.2G113800 | 208 | 28 | 50 | 18 | 0.011 | 0.154 |
| CepurR40.2G168200 | 62 | 0 | 12 | 2 | 0.012 | 0.357 |
| CepurR40.3G237700 | 48 | 21 | 8 | 1 | 0.012 | 0.423 |
| CepurR40.1G126200 | 10 | 1 | 4 | 0 | 0.012 | 0.722 |
| CepurR40.6G072600 | 11 | 2 | 0 | 3 | 0.012 | -0.846 |
| CepurR40.1G014500 | 5 | 0 | 5 | 1 | 0.013 | 0.606 |
| CepurR40.10G192200 | 10 | 0 | 1 | 5 | 0.013 | -0.667 |
| CepurR40.1G000600 | 11 | 0 | 0 | 2 | 0.013 | -1 |
| CepurR40.5G146600 | 32 | 0 | 0 | 10 | 0.013 | -0.39 |
| CepurR40.7G079600 | 17 | 6 | 0 | 5 | 0.014 | -0.63 |
| CepurR40.5G105100 | 80 | 10 | 10 | 0 | 0.014 | 0.407 |
| CepurR40.2G071700 | 32 | 0 | 10 | 13 | 0.014 | -0.327 |
| CepurR40.11G044500 | 19 | 1 | 3 | 0 | 0.014 | 0.782 |
| CepurR40.5G136800 | 7 | 2 | 0 | 14 | 0.014 | -0.318 |
| CepurR40.4G032700 | 92 | 1 | 3 | 5 | 0.014 | -0.432 |
| CepurR40.11G019100 | 12 | 1 | 6 | 1 | 0.015 | 0.541 |
| CepurR40.4G129300 | 50 | 5 | 12 | 1 | 0.015 | 0.355 |
| CepurR40.4G253000 | 8 | 0 | 3 | 12 | 0.015 | -0.527 |
| CepurR40.4G168500 | 36 | 2 | 0 | 4 | 0.015 | -0.667 |
| CepurR40.6G024700 | 106 | 0 | 23 | 24 | 0.015 | -0.199 |
| CepurR40.5G103800 | 17 | 2 | 15 | 11 | 0.015 | 0.311 |
| CepurR40.9G008700 | 10 | 0 | 4 | 16 | 0.016 | -0.467 |
| CepurR40.2G118600 | 27 | 9 | 3 | 0 | 0.016 | 0.745 |
| CepurR40.5G014100 | 30 | 5 | 0 | 3 | 0.017 | -0.811 |
| CepurR40.11G134500 | 30 | 2 | 5 | 0 | 0.017 | 0.552 |
| CepurR40.9G104500 | 21 | 0 | 1 | 4 | 0.017 | -0.608 |
| CepurR40.2G116000 | 58 | 8 | 0 | 3 | 0.017 | -0.744 |
| CepurR40.2G281300 | 30 | 8 | 4 | 0 | 0.017 | 0.651 |
| CepurR40.10G143200 | 18 | 4 | 4 | 7 | 0.017 | -0.584 |
| CepurR40.4G200300 | 28 | 3 | 0 | 3 | 0.018 | -0.757 |

| | | | | | | |
|---|---|---|---|---|---|---|
| CepurR40.5G198800 | 0 | 0 | 3 | 0 | 0.018 | 1 |
| CepurR40.7G151300 | 23 | 0 | 14 | 11 | 0.018 | 0.269 |
| CepurR40.9G006300 | 42 | 1 | 2 | 12 | 0.018 | -0.351 |
| CepurR40.4G177400 | 9 | 0 | 10 | 3 | 0.019 | 0.409 |
| CepurR40.11G086200 | 39 | 3 | 5 | 8 | 0.019 | -0.38 |
| CepurR40.1G286200 | 13 | 0 | 1 | 9 | 0.019 | -0.491 |
| CepurR40.4G075200 | 9 | 0 | 0 | 4 | 0.019 | -0.818 |
| CepurR40.10G140100 | 6 | 0 | 4 | 0 | 0.02 | 0.684 |
| CepurR40.1G284400 | 18 | 2 | 16 | 6 | 0.02 | 0.352 |
| CepurR40.6G196400 | 6 | 1 | 7 | 4 | 0.02 | 0.422 |
| CepurR40.11G075700 | 8 | 2 | 4 | 1 | 0.021 | 0.605 |
| CepurR40.2G235700 | 110 | 30 | 2 | 5 | 0.021 | -0.438 |
| CepurR40.3G029700 | 10 | 1 | 2 | 10 | 0.022 | -0.458 |
| CepurR40.4G234300 | 20 | 2 | 1 | 7 | 0.022 | -0.463 |
| CepurR40.7G058300 | 5 | 1 | 10 | 6 | 0.022 | 0.408 |
| CepurR40.7G050100 | 2 | 0 | 3 | 0 | 0.022 | 0.833 |
| CepurR40.2G216600 | 36 | 0 | 8 | 0 | 0.022 | 0.419 |
| CepurR40.11G028900 | 14 | 1 | 0 | 4 | 0.022 | -0.667 |
| CepurR40.7G011900 | 0 | 0 | 2 | 0 | 0.022 | 1 |
| CepurR40.2G134400 | 1 | 0 | 3 | 1 | 0.022 | 0.673 |
| CepurR40.11G159200 | 22 | 3 | 0 | 5 | 0.022 | -0.564 |
| CepurR40.7G109000 | 35 | 3 | 4 | 8 | 0.022 | -0.353 |
| CepurR40.8G192400 | 57 | 0 | 2 | 5 | 0.022 | -0.455 |
| CepurR40.5G188700 | 45 | 4 | 8 | 17 | 0.023 | -0.314 |
| CepurR40.4G176400 | 12 | 1 | 4 | 14 | 0.023 | -0.378 |
| CepurR40.4G245200 | 143 | 2 | 2 | 5 | 0.023 | -0.437 |
| CepurR40.8G170800 | 23 | 7 | 0 | 9 | 0.023 | -0.39 |
| CepurR40.2G099100 | 21 | 0 | 6 | 0 | 0.023 | 0.543 |
| CepurR40.1G117700 | 12 | 0 | 1 | 5 | 0.023 | -0.583 |
| CepurR40.5G104100 | 20 | 2 | 6 | 0 | 0.023 | 0.565 |
| CepurR40.7G026600 | 34 | 16 | 0 | 3 | 0.023 | -0.708 |
| CepurR40.1G271700 | 7 | 4 | 0 | 7 | 0.023 | -0.438 |
| CepurR40.2G133900 | 11 | 2 | 4 | 0 | 0.024 | 0.656 |
| CepurR40.6G067300 | 7 | 3 | 5 | 1 | 0.024 | 0.6 |
| CepurR40.4G113000 | 49 | 13 | 2 | 9 | 0.024 | -0.381 |
| CepurR40.9G030600 | 10 | 0 | 0 | 6 | 0.024 | -0.556 |
| CepurR40.5G118100 | 15 | 2 | 4 | 0 | 0.024 | 0.625 |
| CepurR40.12G100700 | 12 | 0 | 0 | 3 | 0.025 | -0.8 |
| CepurR40.1G110800 | 14 | 0 | 3 | 0 | 0.025 | 0.725 |
| CepurR40.9G063700 | 11 | 1 | 4 | 0 | 0.025 | 0.667 |
| CepurR40.8G127800 | 37 | 2 | 3 | 12 | 0.025 | -0.329 |
| CepurR40.2G240200 | 21 | 6 | 1 | 7 | 0.025 | -0.4 |
| CepurR40.5G104600 | 25 | 2 | 13 | 3 | 0.025 | 0.332 |
| CepurR40.11G022400 | 18 | 0 | 4 | 7 | 0.026 | -0.419 |

| | | | | | | |
|---|---|---|---|---|---|---|
| CepurR40.1G016700 | 25 | 4 | 16 | 0 | 0.026 | 0.242 |
| CepurR40.10G053900 | 2 | 2 | 4 | 2 | 0.026 | 0.524 |
| CepurR40.1G138700 | 10 | 1 | 10 | 0 | 0.027 | 0.286 |
| CepurR40.3G000200 | 49 | 10 | 10 | 1 | 0.027 | 0.365 |
| CepurR40.12G080300 | 6 | 0 | 4 | 1 | 0.027 | 0.539 |
| CepurR40.12G163300 | 2 | 0 | 3 | 0 | 0.027 | 0.778 |
| CepurR40.12G131000 | 0 | 0 | 2 | 0 | 0.028 | 1 |
| CepurR40.9G099000 | 54 | 3 | 6 | 0 | 0.028 | 0.505 |
| CepurR40.7G072300 | 28 | 4 | 0 | 3 | 0.028 | -0.737 |
| CepurR40.2G180200 | 64 | 8 | 0 | 6 | 0.028 | -0.489 |
| CepurR40.3G137200 | 27 | 0 | 1 | 4 | 0.028 | -0.55 |
| CepurR40.10G006300 | 0 | 0 | 3 | 0 | 0.029 | 1 |
| CepurR40.6G002200 | 4 | 0 | 0 | 3 | 0.029 | -1 |
| CepurR40.3G237900 | 14 | 0 | 1 | 6 | 0.029 | -0.557 |
| CepurR40.5G116900 | 15 | 0 | 13 | 20 | 0.029 | 0.213 |
| CepurR40.3G044000 | 7 | 0 | 4 | 1 | 0.029 | 0.55 |
| CepurR40.11G084800 | 6 | 0 | 9 | 3 | 0.029 | 0.434 |
| CepurR40.11G024500 | 14 | 0 | 26 | 11 | 0.029 | 0.265 |
| CepurR40.12G014400 | 26 | 7 | 4 | 9 | 0.029 | -0.359 |
| CepurR40.2G143500 | 22 | 0 | 13 | 5 | 0.029 | 0.315 |
| CepurR40.3G012500 | 3 | 0 | 3 | 0 | 0.029 | 0.786 |
| CepurR40.7G003900 | 9 | 0 | 0 | 4 | 0.029 | -0.692 |
| CepurR40.2G134300 | 29 | 0 | 1 | 8 | 0.029 | -0.416 |
| CepurR40.1G061100 | 0 | 0 | 4 | 9 | 0.03 | 0.308 |
| CepurR40.5G130500 | 29 | 2 | 3 | 10 | 0.031 | -0.338 |
| CepurR40.6G128100 | 25 | 6 | 2 | 10 | 0.031 | -0.314 |
| CepurR40.10G036700 | 13 | 0 | 0 | 3 | 0.031 | -0.765 |
| CepurR40.10G045800 | 91 | 2 | 10 | 13 | 0.031 | -0.26 |
| CepurR40.10G137700 | 43 | 5 | 12 | 0 | 0.031 | 0.283 |
| CepurR40.10G125800 | 18 | 3 | 9 | 1 | 0.032 | 0.426 |
| CepurR40.11G103200 | 16 | 0 | 0 | 2 | 0.032 | -0.889 |
| CepurR40.2G070800 | 16 | 0 | 0 | 2 | 0.032 | -0.889 |
| CepurR40.11G140600 | 52 | 7 | 34 | 12 | 0.032 | 0.219 |
| CepurR40.2G248800 | 48 | 1 | 1 | 4 | 0.032 | -0.538 |
| CepurR40.8G162300 | 16 | 0 | 15 | 6 | 0.032 | 0.304 |
| CepurR40.1G100800 | 8 | 1 | 1 | 23 | 0.032 | -0.292 |
| CepurR40.11G056000 | 16 | 2 | 12 | 3 | 0.032 | 0.356 |
| CepurR40.7G030900 | 48 | 1 | 17 | 0 | 0.032 | 0.238 |
| CepurR40.1G277300 | 7 | 0 | 13 | 7 | 0.032 | 0.346 |
| CepurR40.3G178700 | 29 | 3 | 2 | 4 | 0.033 | -0.495 |
| CepurR40.7G055400 | 17 | 0 | 0 | 6 | 0.033 | -0.5 |
| CepurR40.2G063000 | 13 | 7 | 5 | 6 | 0.033 | -0.412 |
| CepurR40.5G104200 | 10 | 1 | 0 | 2 | 0.033 | -0.909 |
| CepurR40.4G090600 | 118 | 15 | 20 | 21 | 0.033 | -0.179 |

| | | | | | | |
|---|---|---|---|---|---|---|
| CepurR40.7G038700 | 13 | 1 | 2 | 6 | 0.033 | -0.472 |
| CepurR40.11G014100 | 1 | 0 | 3 | 0 | 0.033 | 0.857 |
| CepurR40.7G000900 | 4 | 0 | 7 | 0 | 0.034 | 0.556 |
| CepurR40.2G198900 | 121 | 0 | 15 | 2 | 0.034 | 0.271 |
| CepurR40.6G131400 | 16 | 0 | 7 | 4 | 0.034 | 0.351 |
| CepurR40.6G147700 | 18 | 1 | 9 | 1 | 0.034 | 0.4 |
| CepurR40.1G210300 | 19 | 0 | 6 | 1 | 0.035 | 0.485 |
| CepurR40.6G076900 | 8 | 0 | 0 | 3 | 0.035 | -0.889 |
| CepurR40.7G075500 | 7 | 1 | 0 | 3 | 0.035 | -0.778 |
| CepurR40.7G187800 | 8 | 0 | 0 | 3 | 0.035 | -0.8 |
| CepurR40.11G032800 | 10 | 4 | 1 | 7 | 0.035 | -0.431 |
| CepurR40.7G026300 | 2 | 4 | 0 | 2 | 0.036 | -1 |
| CepurR40.8G133700 | 0 | 0 | 2 | 0 | 0.036 | 1 |
| CepurR40.8G154700 | 21 | 2 | 2 | 6 | 0.036 | -0.45 |
| CepurR40.2G079900 | 21 | 0 | 0 | 6 | 0.036 | -0.477 |
| CepurR40.10G154400 | 12 | 0 | 4 | 3 | 0.036 | -0.429 |
| CepurR40.11G132900 | 15 | 0 | 9 | 2 | 0.036 | 0.377 |
| CepurR40.3G025800 | 14 | 0 | 0 | 3 | 0.036 | -0.737 |
| CepurR40.7G094000 | 11 | 2 | 5 | 3 | 0.037 | 0.405 |
| CepurR40.1G048000 | 9 | 1 | 9 | 3 | 0.037 | 0.429 |
| CepurR40.3G178200 | 6 | 4 | 3 | 0 | 0.037 | 0.739 |
| CepurR40.4G091600 | 12 | 6 | 5 | 9 | 0.037 | -0.31 |
| CepurR40.5G069000 | 28 | 1 | 9 | 1 | 0.037 | 0.381 |
| CepurR40.3G138700 | 27 | 6 | 1 | 4 | 0.037 | -0.53 |
| CepurR40.2G131000 | 22 | 3 | 4 | 1 | 0.037 | 0.511 |
| CepurR40.8G099600 | 7 | 2 | 6 | 1 | 0.037 | 0.539 |
| CepurR40.6G010700 | 12 | 0 | 0 | 4 | 0.037 | -0.632 |
| CepurR40.7G076400 | 7 | 0 | 5 | 0 | 0.037 | 0.611 |
| CepurR40.11G059000 | 7 | 1 | 8 | 5 | 0.037 | 0.382 |
| CepurR40.1G276900 | 12 | 1 | 1 | 3 | 0.037 | -0.607 |
| CepurR40.6G000700 | 9 | 3 | 2 | 4 | 0.037 | -0.485 |
| CepurR40.5G158800 | 20 | 2 | 8 | 0 | 0.038 | 0.459 |
| CepurR40.12G154800 | 4 | 2 | 0 | 7 | 0.038 | -1 |
| CepurR40.3G077300 | 3 | 0 | 6 | 5 | 0.038 | 0.395 |
| CepurR40.3G145700 | 93 | 0 | 8 | 6 | 0.039 | -0.252 |
| CepurR40.1G064900 | 9 | 0 | 4 | 18 | 0.039 | -0.418 |
| CepurR40.5G145300 | 14 | 0 | 0 | 2 | 0.039 | -0.875 |
| CepurR40.1G256900 | 12 | 6 | 6 | 8 | 0.039 | -0.321 |
| CepurR40.1G307000 | 1 | 1 | 13 | 37 | 0.04 | 0.212 |
| CepurR40.7G146100 | 103 | 7 | 1 | 4 | 0.04 | -0.487 |
| CepurR40.8G064200 | 18 | 3 | 6 | 1 | 0.04 | 0.482 |
| CepurR40.11G014400 | 18 | 0 | 9 | 2 | 0.04 | 0.379 |
| CepurR40.10G151200 | 169 | 39 | 11 | 0 | 0.041 | 0.281 |
| CepurR40.11G173300 | 12 | 2 | 8 | 18 | 0.041 | -0.359 |

| | | | | | | |
|---|---|---|---|---|---|---|
| CepurR40.3G229500 | 22 | 0 | 8 | 0 | 0.041 | 0.371 |
| CepurR40.5G055600 | 8 | 0 | 5 | 0 | 0.041 | 0.579 |
| CepurR40.8G086900 | 23 | 2 | 5 | 6 | 0.041 | -0.367 |
| CepurR40.12G109300 | 5 | 0 | 6 | 4 | 0.041 | 0.4 |
| CepurR40.5G111400 | 27 | 2 | 1 | 6 | 0.041 | -0.457 |
| CepurR40.8G196200 | 9 | 2 | 2 | 8 | 0.041 | -0.492 |
| CepurR40.11G016000 | 14 | 0 | 4 | 0 | 0.041 | 0.588 |
| CepurR40.4G016400 | 15 | 0 | 0 | 3 | 0.042 | -0.714 |
| CepurR40.9G116500 | 1 | 0 | 2 | 2 | 0.042 | 0.455 |
| CepurR40.11G148700 | 7 | 2 | 4 | 1 | 0.042 | 0.567 |
| CepurR40.1G311500 | 15 | 0 | 1 | 7 | 0.043 | -0.475 |
| CepurR40.12G129900 | 10 | 3 | 4 | 3 | 0.043 | 0.399 |
| CepurR40.2G208100 | 51 | 6 | 6 | 9 | 0.043 | -0.299 |
| CepurR40.3G180400 | 11 | 0 | 0 | 3 | 0.043 | -0.733 |
| CepurR40.5G085700 | 9 | 2 | 0 | 3 | 0.043 | -0.692 |
| CepurR40.10G110000 | 48 | 2 | 13 | 29 | 0.043 | -0.18 |
| CepurR40.10G058400 | 22 | 2 | 0 | 3 | 0.043 | -0.667 |
| CepurR40.10G074600 | 28 | 4 | 12 | 3 | 0.043 | 0.317 |
| CepurR40.1G082800 | 18 | 4 | 0 | 3 | 0.044 | -0.667 |
| CepurR40.11G086300 | 8 | 0 | 0 | 5 | 0.045 | -0.571 |
| CepurR40.5G168200 | 29 | 0 | 0 | 3 | 0.045 | -0.674 |
| CepurR40.1G021900 | 309 | 85 | 0 | 3 | 0.045 | -0.655 |
| CepurR40.2G226000 | 29 | 1 | 4 | 0 | 0.045 | 0.561 |
| CepurR40.2G000400 | 7 | 2 | 0 | 5 | 0.045 | -0.5 |
| CepurR40.3G099500 | 48 | 18 | 0 | 4 | 0.045 | -0.533 |
| CepurR40.3G268800 | 6 | 1 | 0 | 3 | 0.045 | -0.75 |
| CepurR40.8G113900 | 31 | 0 | 9 | 0 | 0.046 | 0.354 |
| CepurR40.5G074300 | 19 | 1 | 3 | 6 | 0.046 | -0.397 |
| CepurR40.11G071700 | 19 | 1 | 1 | 7 | 0.046 | -0.451 |
| CepurR40.4G247500 | 19 | 2 | 1 | 6 | 0.046 | -0.433 |
| CepurR40.5G060700 | 28 | 6 | 0 | 3 | 0.046 | -0.667 |
| CepurR40.8G115600 | 30 | 3 | 7 | 5 | 0.046 | -0.299 |
| CepurR40.3G083000 | 9 | 0 | 0 | 6 | 0.046 | -0.6 |
| CepurR40.11G098700 | 7 | 0 | 2 | 0 | 0.046 | 0.816 |
| CepurR40.10G034900 | 58 | 1 | 20 | 6 | 0.046 | 0.222 |
| CepurR40.7G039300 | 47 | 3 | 18 | 16 | 0.046 | -0.229 |
| CepurR40.9G143600 | 21 | 2 | 0 | 3 | 0.047 | -0.7 |
| CepurR40.5G007700 | 6 | 3 | 7 | 20 | 0.047 | -0.241 |
| CepurR40.4G007500 | 6 | 0 | 2 | 0 | 0.047 | 0.818 |
| CepurR40.11G168500 | 13 | 2 | 4 | 0 | 0.047 | 0.519 |
| CepurR40.1G194100 | 17 | 3 | 1 | 4 | 0.047 | -0.508 |
| CepurR40.2G169700 | 21 | 26 | 1 | 6 | 0.047 | -0.557 |
| CepurR40.7G133900 | 15 | 1 | 1 | 4 | 0.047 | -0.514 |
| CepurR40.11G034300 | 279 | 16 | 14 | 1 | 0.047 | 0.246 |

| | | | | | | |
|---|---|---|---|---|---|---|
| CepurR40.11G095100 | 51 | 14 | 29 | 8 | 0.047 | 0.198 |
| CepurR40.11G002600 | 5 | 0 | 0 | 2 | 0.048 | -1 |
| CepurR40.12G158500 | 2 | 0 | 0 | 5 | 0.048 | -1 |
| CepurR40.1G140500 | 5 | 0 | 1 | 3 | 0.048 | -0.75 |
| CepurR40.11G053400 | 19 | 5 | 16 | 28 | 0.048 | -0.292 |
| CepurR40.7G057900 | 4 | 2 | 6 | 6 | 0.049 | 0.362 |
| CepurR40.10G108900 | 5 | 2 | 2 | 0 | 0.049 | 0.848 |
| CepurR40.7G031200 | 16 | 7 | 4 | 8 | 0.049 | -0.307 |
| CepurR40.3G078000 | 12 | 0 | 5 | 14 | 0.049 | -0.368 |
| CepurR40.1G316900 | 3 | 2 | 2 | 9 | 0.049 | -0.568 |
| CepurR40.3G055800 | 5 | 0 | 12 | 0 | 0.049 | 0.375 |
| CepurR40.5G156100 | 5 | 0 | 4 | 2 | 0.049 | 0.458 |
| CepurR40.5G166400 | 39 | 0 | 1 | 4 | 0.049 | -0.484 |
| CepurR40.5G042700 | 180 | 30 | 8 | 11 | 0.049 | -0.236 |
| CepurR40.7G071100 | 7 | 1 | 2 | 0 | 0.05 | 0.8 |
| CepurR40.4G241200 | 18 | 1 | 0 | 2 | 0.05 | -0.818 |
| CepurR40.2G042400 | 25 | 0 | 0 | 5 | 0.05 | -0.556 |
| CepurR40.10G138200 | 12 | 1 | 7 | 8 | 0.05 | -0.39 |
| CepurR40.3G048200 | 8 | 0 | 3 | 0 | 0.05 | 0.68 |
| CepurR40.11G141100 | 48 | 22 | 0 | 3 | 0.05 | -0.632 |
| CepurR40.7G088500 | 20 | 0 | 2 | 3 | 0.05 | -0.47 |
| CepurR40.1G123500 | 13 | 0 | 2 | 0 | 0.05 | 0.74 |
| CepurR40.3G162200 | 11 | 8 | 6 | 9 | 0.051 | -0.211 |
| CepurR40.12G038500 | 6 | 0 | 8 | 2 | 0.051 | 0.425 |
| CepurR40.2G263400 | 1 | 0 | 3 | 1 | 0.052 | 0.639 |
| CepurR40.4G043000 | 23 | 0 | 1 | 4 | 0.052 | -0.497 |
| CepurR40.9G050800 | 30 | 10 | 1 | 4 | 0.052 | -0.438 |
| CepurR40.10G179800 | 19 | 9 | 8 | 3 | 0.052 | 0.394 |
| CepurR40.1G021800 | 65 | 1 | 0 | 8 | 0.052 | -0.367 |
| CepurR40.2G051000 | 28 | 1 | 1 | 3 | 0.052 | -0.528 |
| CepurR40.4G188600 | 15 | 0 | 0 | 3 | 0.052 | -0.682 |
| CepurR40.8G113000 | 5 | 0 | 2 | 12 | 0.052 | -0.482 |
| CepurR40.5G136000 | 16 | 2 | 2 | 7 | 0.052 | -0.444 |
| CepurR40.4G255400 | 19 | 1 | 1 | 3 | 0.052 | -0.542 |
| CepurR40.10G116100 | 10 | 1 | 2 | 3 | 0.053 | -0.509 |
| CepurR40.7G034000 | 14 | 1 | 0 | 2 | 0.053 | -0.824 |
| CepurR40.9G028500 | 14 | 1 | 0 | 2 | 0.053 | -0.824 |
| CepurR40.5G160400 | 4 | 0 | 7 | 4 | 0.053 | 0.401 |
| CepurR40.3G004900 | 6 | 0 | 0 | 10 | 0.053 | -0.375 |
| CepurR40.6G150800 | 11 | 3 | 8 | 6 | 0.054 | 0.303 |
| CepurR40.1G075100 | 3 | 1 | 2 | 1 | 0.054 | 0.578 |
| CepurR40.6G170600 | 9 | 0 | 0 | 5 | 0.054 | -0.529 |
| CepurR40.1G124900 | 6 | 2 | 0 | 7 | 0.054 | -0.429 |
| CepurR40.7G112200 | 12 | 16 | 4 | 6 | 0.054 | -0.2 |

| | | | | | | |
|---|---|---|---|---|---|---|
| CepurR40.9G054900 | 68 | 11 | 23 | 10 | 0.054 | 0.228 |
| CepurR40.2G222300 | 55 | 12 | 4 | 7 | 0.055 | -0.315 |
| CepurR40.2G060000 | 6 | 2 | 0 | 2 | 0.055 | -1 |
| CepurR40.4G011600 | 8 | 0 | 0 | 2 | 0.055 | -0.889 |
| CepurR40.8G168600 | 1 | 0 | 2 | 0 | 0.055 | 0.875 |
| CepurR40.5G147800 | 24 | 4 | 3 | 0 | 0.055 | 0.631 |
| CepurR40.2G115600 | 29 | 2 | 0 | 3 | 0.055 | -0.69 |
| CepurR40.6G079400 | 21 | 7 | 0 | 5 | 0.055 | -0.583 |
| CepurR40.2G181400 | 17 | 2 | 7 | 1 | 0.055 | 0.428 |
| CepurR40.7G027600 | 27 | 1 | 2 | 10 | 0.055 | -0.324 |
| CepurR40.5G170000 | 27 | 0 | 2 | 6 | 0.056 | -0.393 |
| CepurR40.6G087000 | 33 | 12 | 2 | 6 | 0.056 | -0.373 |
| CepurR40.7G151200 | 13 | 0 | 7 | 13 | 0.056 | -0.334 |
| CepurR40.2G143800 | 17 | 0 | 4 | 1 | 0.056 | 0.473 |
| CepurR40.6G103200 | 11 | 3 | 1 | 2 | 0.056 | -0.583 |
| CepurR40.5G075000 | 7 | 1 | 3 | 0 | 0.056 | 0.682 |
| CepurR40.3G019000 | 27 | 1 | 5 | 0 | 0.056 | 0.5 |
| CepurR40.5G135200 | 16 | 1 | 0 | 5 | 0.056 | -0.485 |
| CepurR40.4G262900 | 1 | 1 | 6 | 2 | 0.057 | 0.607 |
| CepurR40.12G087100 | 39 | 1 | 1 | 7 | 0.057 | -0.409 |
| CepurR40.11G042500 | 72 | 5 | 3 | 6 | 0.057 | -0.366 |
| CepurR40.5G089200 | 10 | 3 | 0 | 5 | 0.058 | -0.476 |
| CepurR40.11G102700 | 54 | 2 | 13 | 1 | 0.058 | 0.27 |
| CepurR40.6G017000 | 9 | 0 | 6 | 0 | 0.058 | 0.471 |
| CepurR40.4G183900 | 25 | 0 | 10 | 2 | 0.058 | 0.301 |
| CepurR40.9G050100 | 192 | 26 | 7 | 7 | 0.059 | -0.247 |
| CepurR40.7G035900 | 279 | 0 | 1 | 4 | 0.059 | -0.444 |
| CepurR40.2G212500 | 25 | 0 | 0 | 3 | 0.059 | -0.641 |
| CepurR40.3G017900 | 4 | 0 | 3 | 1 | 0.059 | 0.483 |
| CepurR40.4G033000 | 25 | 0 | 4 | 9 | 0.06 | -0.317 |
| CepurR40.2G177600 | 51 | 3 | 4 | 3 | 0.06 | -0.308 |
| CepurR40.10G114100 | 10 | 1 | 0 | 5 | 0.06 | -0.526 |
| CepurR40.11G147000 | 10 | 1 | 5 | 0 | 0.06 | 0.5 |
| CepurR40.2G177700 | 84 | 10 | 5 | 0 | 0.06 | 0.5 |
| CepurR40.4G161000 | 5 | 6 | 1 | 11 | 0.06 | -0.211 |
| CepurR40.4G162400 | 3 | 0 | 2 | 1 | 0.06 | 0.551 |
| CepurR40.8G089900 | 27 | 5 | 8 | 18 | 0.06 | -0.232 |
| CepurR40.2G129300 | 14 | 0 | 1 | 5 | 0.06 | -0.5 |
| CepurR40.10G132600 | 16 | 1 | 0 | 3 | 0.06 | -0.667 |
| CepurR40.3G057500 | 5 | 4 | 3 | 0 | 0.06 | 0.688 |
| CepurR40.9G062300 | 15 | 8 | 0 | 2 | 0.06 | -0.833 |
| CepurR40.2G069300 | 36 | 7 | 2 | 10 | 0.06 | -0.289 |
| CepurR40.2G175900 | 29 | 0 | 3 | 0 | 0.06 | 0.623 |
| CepurR40.1G299300 | 9 | 2 | 3 | 0 | 0.061 | 0.679 |

| | | | | | | |
|---|---|---|---|---|---|---|
| CepurR40.3G253600 | 11 | 0 | 3 | 1 | 0.061 | 0.511 |
| CepurR40.12G142200 | 21 | 1 | 5 | 4 | 0.061 | -0.357 |
| CepurR40.3G157600 | 11 | 2 | 1 | 10 | 0.061 | -0.332 |
| CepurR40.5G124100 | 214 | 4 | 15 | 14 | 0.061 | -0.173 |
| CepurR40.12G127400 | 42 | 13 | 2 | 6 | 0.062 | -0.35 |
| CepurR40.2G125600 | 32 | 1 | 5 | 0 | 0.062 | 0.484 |
| CepurR40.5G167100 | 12 | 0 | 6 | 0 | 0.062 | 0.429 |
| CepurR40.4G254700 | 4 | 2 | 9 | 12 | 0.062 | 0.269 |
| CepurR40.9G137800 | 32 | 3 | 0 | 3 | 0.062 | -0.604 |
| CepurR40.10G126000 | 10 | 0 | 2 | 5 | 0.062 | -0.484 |
| CepurR40.11G092500 | 15 | 0 | 0 | 1 | 0.063 | -1 |
| CepurR40.4G213700 | 21 | 0 | 1 | 3 | 0.063 | -0.625 |
| CepurR40.11G102400 | 132 | 16 | 9 | 12 | 0.063 | -0.235 |
| CepurR40.9G062500 | 81 | 15 | 1 | 4 | 0.063 | -0.438 |
| CepurR40.5G182200 | 23 | 3 | 14 | 2 | 0.063 | 0.27 |
| CepurR40.11G006800 | 10 | 0 | 1 | 3 | 0.063 | -0.583 |
| CepurR40.7G047100 | 15 | 3 | 0 | 3 | 0.064 | -0.625 |
| CepurR40.9G066700 | 10 | 2 | 3 | 0 | 0.064 | 0.667 |
| CepurR40.8G073600 | 12 | 5 | 2 | 4 | 0.064 | -0.524 |
| CepurR40.11G103300 | 24 | 0 | 0 | 3 | 0.064 | -0.632 |
| CepurR40.9G175700 | 130 | 0 | 3 | 6 | 0.064 | -0.344 |
| CepurR40.4G241700 | 13 | 1 | 0 | 6 | 0.065 | -0.433 |
| CepurR40.5G135600 | 5 | 0 | 2 | 0 | 0.065 | 0.792 |
| CepurR40.4G066100 | 6 | 3 | 3 | 6 | 0.065 | -0.417 |
| CepurR40.2G075000 | 11 | 2 | 0 | 2 | 0.065 | -0.786 |
| CepurR40.4G159300 | 15 | 2 | 3 | 0 | 0.066 | 0.643 |
| CepurR40.7G067500 | 12 | 6 | 5 | 0 | 0.066 | 0.478 |
| CepurR40.3G126700 | 49 | 4 | 0 | 3 | 0.066 | -0.62 |
| CepurR40.2G123700 | 8 | 2 | 6 | 0 | 0.066 | 0.467 |
| CepurR40.1G026800 | 5 | 2 | 4 | 18 | 0.067 | -0.235 |
| CepurR40.11G034600 | 0 | 0 | 2 | 0 | 0.067 | 1 |
| CepurR40.11G098400 | 7 | 0 | 0 | 2 | 0.067 | -0.875 |
| CepurR40.12G091500 | 0 | 0 | 4 | 0 | 0.067 | 1 |
| CepurR40.1G009500 | 4 | 0 | 0 | 2 | 0.067 | -1 |
| CepurR40.3G020100 | 3 | 1 | 0 | 2 | 0.067 | -1 |
| CepurR40.4G106800 | 7 | 0 | 0 | 2 | 0.067 | -0.875 |
| CepurR40.4G251200 | 6 | 1 | 1 | 2 | 0.067 | -0.667 |
| CepurR40.6G089000 | 5 | 2 | 1 | 2 | 0.067 | -0.667 |
| CepurR40.7G113000 | 7 | 0 | 1 | 2 | 0.067 | -0.667 |
| CepurR40.7G191300 | 0 | 0 | 2 | 0 | 0.067 | 1 |
| CepurR40.11G057900 | 5 | 2 | 1 | 8 | 0.067 | -0.389 |
| CepurR40.7G093700 | 8 | 0 | 2 | 0 | 0.068 | 0.765 |
| CepurR40.11G037500 | 5 | 1 | 6 | 5 | 0.068 | 0.328 |
| CepurR40.1G018600 | 15 | 1 | 16 | 7 | 0.068 | 0.227 |

| | | | | | | |
|---|---|---|---|---|---|---|
| CepurR40.1G106400 | 2 | 0 | 1 | 0 | 0.068 | 0.952 |
| CepurR40.3G012200 | 5 | 0 | 4 | 3 | 0.068 | 0.379 |
| CepurR40.2G062700 | 24 | 2 | 0 | 5 | 0.068 | -0.453 |
| CepurR40.7G189500 | 4 | 6 | 0 | 3 | 0.069 | -0.5 |
| CepurR40.8G108500 | 9 | 1 | 0 | 3 | 0.069 | -0.643 |
| CepurR40.11G033200 | 8 | 2 | 1 | 6 | 0.069 | -0.657 |
| CepurR40.7G049700 | 35 | 0 | 5 | 0 | 0.069 | 0.444 |
| CepurR40.11G068000 | 24 | 0 | 2 | 7 | 0.069 | -0.426 |
| CepurR40.5G024100 | 8 | 0 | 3 | 6 | 0.07 | -0.467 |
| CepurR40.7G114900 | 3 | 0 | 3 | 0 | 0.07 | 0.7 |
| CepurR40.8G143900 | 6 | 0 | 0 | 4 | 0.07 | -0.667 |
| CepurR40.6G056200 | 40 | 1 | 3 | 0 | 0.071 | 0.583 |
| CepurR40.2G092000 | 60 | 3 | 6 | 6 | 0.071 | -0.279 |
| CepurR40.11G124600 | 81 | 44 | 12 | 13 | 0.071 | -0.184 |
| CepurR40.4G188800 | 236 | 64 | 6 | 5 | 0.071 | -0.257 |
| CepurR40.6G178400 | 14 | 2 | 0 | 5 | 0.071 | -0.452 |
| CepurR40.12G043200 | 31 | 2 | 14 | 5 | 0.071 | 0.237 |
| CepurR40.11G041500 | 14 | 0 | 0 | 3 | 0.072 | -0.636 |
| CepurR40.11G127300 | 21 | 2 | 0 | 2 | 0.073 | -0.808 |
| CepurR40.2G085700 | 4 | 49 | 0 | 3 | 0.073 | -0.222 |
| CepurR40.2G061900 | 1 | 0 | 12 | 7 | 0.073 | 0.465 |
| CepurR40.6G207000 | 8 | 1 | 0 | 8 | 0.073 | -0.364 |
| CepurR40.1G020300 | 3 | 0 | 2 | 1 | 0.073 | 0.542 |
| CepurR40.7G132000 | 40 | 2 | 10 | 3 | 0.073 | 0.293 |
| CepurR40.2G124400 | 28 | 0 | 7 | 2 | 0.073 | 0.354 |
| CepurR40.3G158700 | 15 | 6 | 5 | 21 | 0.073 | -0.192 |
| CepurR40.1G064200 | 5 | 0 | 2 | 1 | 0.074 | 0.532 |
| CepurR40.4G159000 | 63 | 5 | 10 | 2 | 0.074 | 0.286 |
| CepurR40.5G122500 | 2 | 1 | 5 | 2 | 0.074 | 0.464 |
| CepurR40.2G099000 | 29 | 0 | 1 | 5 | 0.074 | -0.45 |
| CepurR40.6G011300 | 7 | 3 | 4 | 6 | 0.074 | -0.475 |
| CepurR40.1G263700 | 18 | 1 | 7 | 0 | 0.075 | 0.419 |
| CepurR40.2G148800 | 12 | 1 | 4 | 8 | 0.075 | -0.373 |
| CepurR40.3G033200 | 3 | 2 | 0 | 5 | 0.075 | -0.5 |
| CepurR40.5G113400 | 9 | 3 | 10 | 2 | 0.076 | 0.333 |
| CepurR40.5G041200 | 0 | 0 | 4 | 2 | 0.076 | 0.667 |
| CepurR40.5G108700 | 0 | 0 | 4 | 2 | 0.076 | 0.667 |
| CepurR40.10G118300 | 20 | 4 | 6 | 3 | 0.076 | 0.344 |
| CepurR40.2G220000 | 265 | 60 | 1 | 3 | 0.076 | -0.468 |
| CepurR40.12G159300 | 16 | 4 | 92 | 54 | 0.077 | 0.173 |
| CepurR40.10G129900 | 12 | 0 | 0 | 1 | 0.077 | -1 |
| CepurR40.1G011000 | 2 | 0 | 2 | 0 | 0.077 | 0.818 |
| CepurR40.1G075400 | 9 | 0 | 1 | 3 | 0.077 | -0.568 |
| CepurR40.1G221700 | 5 | 4 | 2 | 2 | 0.077 | -0.5 |

| | | | | | | |
|---|---|---|---|---|---|---|
| CepurR40.3G005600 | 9 | 0 | 1 | 3 | 0.077 | -0.568 |
| CepurR40.6G019200 | 4 | 0 | 3 | 0 | 0.077 | 0.667 |
| CepurR40.6G088700 | 2 | 0 | 2 | 9 | 0.077 | -0.818 |
| CepurR40.7G188800 | 4 | 0 | 0 | 7 | 0.077 | -0.571 |
| CepurR40.5G128400 | 5 | 0 | 11 | 15 | 0.077 | 0.231 |
| CepurR40.2G101700 | 28 | 1 | 0 | 5 | 0.077 | -0.431 |
| CepurR40.7G099300 | 13 | 0 | 5 | 1 | 0.077 | 0.439 |
| CepurR40.5G059300 | 25 | 5 | 5 | 1 | 0.078 | 0.423 |
| CepurR40.5G069200 | 112 | 3 | 5 | 0 | 0.078 | 0.429 |
| CepurR40.5G104300 | 8 | 2 | 7 | 3 | 0.078 | 0.352 |
| CepurR40.6G184900 | 31 | 1 | 5 | 1 | 0.078 | 0.455 |
| CepurR40.11G158500 | 6 | 3 | 2 | 0 | 0.078 | 0.739 |
| CepurR40.3G154800 | 21 | 0 | 1 | 2 | 0.079 | -0.542 |
| CepurR40.2G078700 | 134 | 0 | 14 | 13 | 0.079 | -0.183 |
| CepurR40.6G132500 | 17 | 7 | 0 | 6 | 0.079 | -0.378 |
| CepurR40.4G133600 | 50 | 10 | 3 | 4 | 0.079 | -0.318 |
| CepurR40.2G225000 | 17 | 2 | 0 | 2 | 0.08 | -0.739 |
| CepurR40.6G129200 | 9 | 4 | 4 | 9 | 0.08 | -0.292 |
| CepurR40.8G196500 | 8 | 4 | 7 | 10 | 0.08 | -0.316 |
| CepurR40.6G062300 | 46 | 17 | 15 | 4 | 0.08 | 0.242 |
| CepurR40.1G299600 | 2 | 0 | 2 | 4 | 0.08 | 0.282 |
| CepurR40.4G222300 | 10 | 7 | 6 | 27 | 0.08 | -0.218 |
| CepurR40.10G067400 | 34 | 0 | 1 | 5 | 0.08 | -0.43 |
| CepurR40.10G000800 | 2 | 3 | 1 | 5 | 0.08 | -0.5 |
| CepurR40.5G127200 | 22 | 4 | 13 | 1 | 0.08 | 0.317 |
| CepurR40.1G330000 | 10 | 0 | 3 | 1 | 0.08 | 0.487 |
| CepurR40.1G034700 | 9 | 0 | 2 | 7 | 0.08 | -0.47 |
| CepurR40.1G089900 | 2 | 3 | 0 | 4 | 0.081 | -1 |
| CepurR40.5G183300 | 4 | 1 | 0 | 4 | 0.081 | -0.571 |
| CepurR40.6G014100 | 5 | 0 | 0 | 4 | 0.081 | -0.625 |
| CepurR40.9G144400 | 89 | 4 | 0 | 3 | 0.081 | -0.578 |
| CepurR40.5G107300 | 243 | 5 | 39 | 16 | 0.081 | 0.129 |
| CepurR40.6G040700 | 1 | 0 | 11 | 1 | 0.081 | 0.417 |
| CepurR40.5G020500 | 260 | 56 | 7 | 8 | 0.082 | -0.218 |
| CepurR40.9G112000 | 51 | 5 | 0 | 3 | 0.082 | -0.567 |
| CepurR40.7G127600 | 57 | 10 | 6 | 0 | 0.082 | 0.412 |
| CepurR40.11G124900 | 3 | 2 | 3 | 0 | 0.082 | 0.75 |
| CepurR40.3G006600 | 4 | 1 | 3 | 0 | 0.082 | 0.692 |
| CepurR40.4G238300 | 8 | 0 | 0 | 4 | 0.082 | -0.615 |
| CepurR40.9G124000 | 38 | 13 | 8 | 12 | 0.082 | -0.244 |
| CepurR40.1G050500 | 19 | 0 | 2 | 3 | 0.082 | -0.426 |
| CepurR40.2G234200 | 13 | 6 | 4 | 3 | 0.082 | -0.295 |
| CepurR40.4G189200 | 52 | 18 | 0 | 3 | 0.083 | -0.536 |
| CepurR40.5G183500 | 8 | 1 | 2 | 0 | 0.083 | 0.714 |

| | | | | | | |
|---|---|---|---|---|---|---|
| CepurR40.2G009300 | 6 | 0 | 0 | 2 | 0.083 | -0.857 |
| CepurR40.4G048300 | 11 | 0 | 0 | 2 | 0.083 | -0.786 |
| CepurR40.4G065500 | 0 | 0 | 2 | 1 | 0.083 | 0.667 |
| CepurR40.4G200100 | 6 | 0 | 0 | 2 | 0.083 | -1 |
| CepurR40.5G062500 | 6 | 0 | 0 | 2 | 0.083 | -0.857 |
| CepurR40.6G115000 | 5 | 1 | 0 | 2 | 0.083 | -0.833 |
| CepurR40.7G057800 | 4 | 2 | 0 | 2 | 0.083 | -0.8 |
| CepurR40.5G022200 | 197 | 30 | 24 | 4 | 0.083 | 0.159 |
| CepurR40.1G294300 | 7 | 2 | 3 | 7 | 0.084 | -0.4 |
| CepurR40.7G123700 | 9 | 0 | 6 | 1 | 0.084 | 0.429 |
| CepurR40.11G095700 | 18 | 2 | 1 | 2 | 0.085 | -0.567 |
| CepurR40.5G023800 | 20 | 0 | 4 | 5 | 0.085 | -0.356 |
| CepurR40.1G293200 | 4 | 0 | 6 | 0 | 0.085 | 0.5 |
| CepurR40.10G136400 | 10 | 4 | 16 | 18 | 0.085 | -0.244 |
| CepurR40.9G123400 | 26 | 6 | 1 | 5 | 0.085 | -0.364 |
| CepurR40.9G090100 | 7 | 1 | 1 | 7 | 0.086 | -0.375 |
| CepurR40.12G090800 | 6 | 0 | 2 | 0 | 0.086 | 0.727 |
| CepurR40.7G175000 | 4 | 2 | 4 | 0 | 0.087 | 0.692 |
| CepurR40.8G101200 | 7 | 2 | 0 | 4 | 0.087 | -0.7 |
| CepurR40.10G134500 | 30 | 7 | 0 | 2 | 0.087 | -0.682 |
| CepurR40.9G027200 | 16 | 5 | 3 | 1 | 0.087 | 0.488 |
| CepurR40.2G211700 | 191 | 39 | 16 | 3 | 0.087 | 0.208 |
| CepurR40.3G128600 | 10 | 0 | 3 | 0 | 0.087 | 0.583 |
| CepurR40.5G117900 | 9 | 1 | 3 | 0 | 0.087 | 0.571 |
| CepurR40.8G062600 | 10 | 0 | 3 | 0 | 0.087 | 0.6 |
| CepurR40.6G020200 | 16 | 2 | 3 | 0 | 0.088 | 0.579 |
| CepurR40.12G046200 | 14 | 3 | 1 | 6 | 0.088 | -0.417 |
| CepurR40.11G131800 | 12 | 1 | 4 | 3 | 0.088 | -0.352 |
| CepurR40.8G065200 | 8 | 5 | 0 | 2 | 0.088 | -0.727 |
| CepurR40.2G084900 | 23 | 1 | 1 | 2 | 0.088 | -0.519 |
| CepurR40.5G162400 | 10 | 1 | 1 | 7 | 0.088 | -0.401 |
| CepurR40.4G269800 | 5 | 0 | 7 | 0 | 0.088 | 0.444 |
| CepurR40.5G184000 | 5 | 0 | 4 | 0 | 0.088 | 0.583 |
| CepurR40.11G045000 | 176 | 14 | 7 | 8 | 0.089 | -0.229 |
| CepurR40.11G148800 | 20 | 7 | 3 | 0 | 0.089 | 0.623 |
| CepurR40.3G236400 | 8 | 0 | 1 | 4 | 0.089 | -0.6 |
| CepurR40.7G190400 | 2 | 0 | 4 | 1 | 0.089 | 0.6 |
| CepurR40.11G028300 | 18 | 1 | 3 | 0 | 0.089 | 0.571 |
| CepurR40.2G188800 | 23 | 0 | 2 | 5 | 0.089 | -0.391 |
| CepurR40.5G120300 | 5 | 1 | 2 | 10 | 0.089 | -0.333 |
| CepurR40.2G269800 | 33 | 0 | 3 | 7 | 0.09 | -0.311 |
| CepurR40.3G242800 | 5 | 8 | 6 | 1 | 0.09 | 0.579 |
| CepurR40.2G107800 | 22 | 1 | 1 | 5 | 0.09 | -0.462 |
| CepurR40.2G086100 | 40 | 1 | 3 | 0 | 0.09 | 0.57 |

| | | | | | | |
|---|---|---|---|---|---|---|
| CepurR40.4G042100 | 7 | 5 | 6 | 1 | 0.09 | 0.42 |
| CepurR40.2G288000 | 7 | 0 | 3 | 0 | 0.09 | 0.611 |
| CepurR40.3G203100 | 6 | 1 | 3 | 0 | 0.09 | 0.647 |
| CepurR40.6G219100 | 4 | 3 | 11 | 0 | 0.09 | 0.333 |
| CepurR40.1G177600 | 62 | 8 | 3 | 2 | 0.09 | -0.299 |
| CepurR40.6G140600 | 9 | 0 | 3 | 1 | 0.091 | 0.477 |
| CepurR40.2G110700 | 53 | 3 | 5 | 10 | 0.091 | -0.256 |
| CepurR40.11G138600 | 5 | 0 | 2 | 0 | 0.091 | 0.737 |
| CepurR40.2G078800 | 15 | 0 | 0 | 2 | 0.091 | -0.75 |
| CepurR40.5G047400 | 5 | 0 | 2 | 0 | 0.091 | 0.75 |
| CepurR40.6G060200 | 9 | 1 | 0 | 1 | 0.091 | -1 |
| CepurR40.7G141700 | 8 | 0 | 2 | 2 | 0.091 | -0.5 |
| CepurR40.1G021300 | 21 | 1 | 5 | 1 | 0.091 | 0.413 |
| CepurR40.2G260300 | 28 | 6 | 1 | 5 | 0.091 | -0.47 |
| CepurR40.2G121700 | 10 | 0 | 13 | 9 | 0.092 | 0.234 |
| CepurR40.1G045700 | 12 | 0 | 3 | 0 | 0.092 | 0.556 |
| CepurR40.8G143600 | 19 | 2 | 2 | 2 | 0.092 | -0.405 |
| CepurR40.10G026000 | 42 | 0 | 8 | 2 | 0.092 | 0.323 |
| CepurR40.3G172700 | 23 | 3 | 2 | 8 | 0.092 | -0.361 |
| CepurR40.8G067800 | 32 | 0 | 7 | 3 | 0.092 | 0.29 |
| CepurR40.10G067300 | 24 | 1 | 3 | 5 | 0.092 | -0.352 |
| CepurR40.2G108500 | 29 | 14 | 5 | 1 | 0.093 | 0.475 |
| CepurR40.6G004900 | 11 | 1 | 1 | 6 | 0.093 | -0.545 |
| CepurR40.1G158300 | 9 | 1 | 2 | 1 | 0.093 | 0.509 |
| CepurR40.1G244000 | 10 | 2 | 0 | 3 | 0.093 | -0.556 |
| CepurR40.3G219800 | 176 | 13 | 1 | 3 | 0.093 | -0.44 |
| CepurR40.3G066000 | 11 | 6 | 1 | 3 | 0.093 | -0.397 |
| CepurR40.11G161500 | 10 | 0 | 1 | 2 | 0.093 | -0.576 |
| CepurR40.7G128300 | 1 | 0 | 2 | 1 | 0.093 | 0.567 |
| CepurR40.7G151500 | 29 | 1 | 5 | 4 | 0.093 | -0.273 |
| CepurR40.1G053800 | 13 | 0 | 8 | 22 | 0.094 | -0.418 |
| CepurR40.7G158900 | 24 | 3 | 2 | 2 | 0.094 | -0.389 |
| CepurR40.11G136600 | 103 | 0 | 12 | 15 | 0.094 | -0.18 |
| CepurR40.8G149300 | 16 | 0 | 5 | 1 | 0.095 | 0.401 |
| CepurR40.5G142100 | 7 | 1 | 2 | 3 | 0.095 | -0.475 |
| CepurR40.6G222200 | 2 | 0 | 3 | 1 | 0.095 | 0.464 |
| CepurR40.11G015700 | 23 | 0 | 11 | 6 | 0.095 | 0.264 |
| CepurR40.4G184100 | 72 | 10 | 6 | 14 | 0.095 | -0.226 |
| CepurR40.10G188900 | 9 | 1 | 0 | 2 | 0.095 | -0.818 |
| CepurR40.1G185500 | 3 | 0 | 2 | 0 | 0.095 | 0.769 |
| CepurR40.3G162700 | 9 | 1 | 0 | 2 | 0.095 | -0.818 |
| CepurR40.6G103300 | 0 | 0 | 2 | 3 | 0.095 | 0.4 |
| CepurR40.6G200000 | 26 | 17 | 1 | 3 | 0.095 | -0.453 |
| CepurR40.1G154300 | 7 | 0 | 1 | 12 | 0.095 | -0.312 |

| | | | | | | |
|---|---|---|---|---|---|---|
| CepurR40.7G021800 | 12 | 0 | 9 | 4 | 0.096 | -0.308 |
| CepurR40.7G139800 | 10 | 0 | 10 | 4 | 0.096 | 0.298 |
| CepurR40.2G199600 | 26 | 0 | 5 | 2 | 0.096 | 0.372 |
| CepurR40.6G049100 | 46 | 4 | 23 | 18 | 0.097 | -0.181 |
| CepurR40.2G187200 | 36 | 5 | 7 | 10 | 0.097 | -0.209 |
| CepurR40.8G032300 | 11 | 0 | 0 | 11 | 0.098 | -0.244 |
| CepurR40.1G174200 | 8 | 0 | 1 | 9 | 0.098 | -0.4 |
| CepurR40.4G146500 | 12 | 0 | 0 | 2 | 0.098 | -0.75 |
| CepurR40.5G191300 | 6 | 3 | 3 | 1 | 0.098 | 0.543 |
| CepurR40.8G084400 | 4 | 0 | 2 | 0 | 0.098 | 0.75 |
| CepurR40.9G058200 | 12 | 0 | 0 | 2 | 0.098 | -0.75 |
| CepurR40.3G167300 | 10 | 1 | 0 | 4 | 0.098 | -0.526 |
| CepurR40.3G240800 | 7 | 4 | 0 | 4 | 0.098 | -0.583 |
| CepurR40.12G044000 | 64 | 10 | 2 | 5 | 0.098 | -0.395 |
| CepurR40.2G057800 | 120 | 26 | 6 | 0 | 0.099 | 0.348 |
| CepurR40.5G116500 | 30 | 1 | 1 | 5 | 0.099 | -0.433 |
| CepurR40.3G159800 | 9 | 1 | 1 | 3 | 0.099 | -0.5 |
| CepurR40.5G052200 | 3 | 0 | 3 | 1 | 0.099 | 0.519 |
| CepurR40.1G023900 | 45 | 6 | 7 | 3 | 0.099 | 0.254 |
| CepurR40.5G055400 | 30 | 8 | 0 | 2 | 0.099 | -0.714 |
| CepurR40.10G166700 | 10 | 1 | 4 | 0 | 0.1 | 0.565 |
| CepurR40.2G133000 | 11 | 0 | 3 | 0 | 0.1 | 0.577 |
| CepurR40.5G173800 | 10 | 1 | 3 | 0 | 0.1 | 0.545 |
| CepurR40.9G055600 | 154 | 24 | 14 | 2 | 0.1 | 0.211 |

**Table S4. Nonsynonymous (*dN*) and synonymous (*dS*) changes on UV genes.** The R40 (male) and GG1 (female) genes analyzed were one-to-one UV orthologs identified in (Carey et al., 2020). Some of the values shown to be zeros did have some changes identified on their branches, but rounding makes them appear zero, so we have indicated these with an asterisk (*).

| R40 gene | R40 *dN* | R40 *dS* | GG1 gene | GG1 *dN* | GG1 *dS* |
|---|---|---|---|---|---|
| CepurR40.VG002900 | 0.032 | 0.178 | CepurGG1.UG204600 | 0.042 | 0.158 |
| CepurR40.VG010000 | 0.002 | 0.022 | CepurGG1.UG274400 | 0.002 | 0.006 |
| CepurR40.VG010200 | 0.004 | 0.019 | CepurGG1.UG059300 | 0.004 | 0.011 |
| CepurR40.VG010300 | 0.018 | 0.164 | CepurGG1.UG337300 | 0.025 | 0.074 |
| CepurR40.VG010700 | 0.041 | 0.152 | CepurGG1.UG270000 | 0.049 | 0.107 |
| CepurR40.VG011000 | 0.019 | 0.122 | CepurGG1.UG207900 | 0.022 | 0.102 |
| CepurR40.VG011600 | 0.021 | 0.077 | CepurGG1.UG016100 | 0.018 | 0.073 |
| CepurR40.VG011900 | 0.019 | 0.079 | CepurGG1.UG338300 | 0.033 | 0.079 |
| CepurR40.VG014300 | 0.022 | 0.099 | CepurGG1.UG317400 | 0.021 | 0.095 |
| CepurR40.VG014800 | 0.004 | 0.255 | CepurGG1.UG197600 | 0.006 | 0.159 |
| CepurR40.VG015400 | 0.03 | 0.059 | CepurGG1.UG291100 | 0.012 | 0.06 |
| CepurR40.VG015600 | 0.005 | 0.093 | CepurGG1.UG259700 | 0.007 | 0.078 |
| CepurR40.VG016500 | 0.033 | 0.149 | CepurGG1.UG208500 | 0.021 | 0.094 |
| CepurR40.VG018900 | 0.004 | 0.027 | CepurGG1.UG005700 | 0* | 0.202 |
| CepurR40.VG020300 | 0.004 | 0.009 | CepurGG1.UG004400 | 0.003 | 0.003 |
| CepurR40.VG021700 | 0.011 | 0.169 | CepurGG1.UG055700 | 0.005 | 0.243 |
| CepurR40.VG022100 | 0.074 | 0.217 | CepurGG1.UG247700 | 0.057 | 0.149 |
| CepurR40.VG022600 | 0.002 | 0.021 | CepurGG1.UG102700 | 0 | 0.002 |
| CepurR40.VG022700 | 0* | 0.085 | CepurGG1.UG002200 | 0 | 0* |
| CepurR40.VG023400 | 0.073 | 0.191 | CepurGG1.UG318500 | 0.054 | 0.175 |
| CepurR40.VG023800 | 0.004 | 0.117 | CepurGG1.UG031000 | 0.001 | 0.125 |
| CepurR40.VG024500 | 0.029 | 0.182 | CepurGG1.UG164900 | 0.014 | 0.097 |
| CepurR40.VG025000 | 0.014 | 0.16 | CepurGG1.UG071300 | 0.016 | 0.052 |
| CepurR40.VG026200 | 0.125 | 0.291 | CepurGG1.UG068700 | 0.089 | 0.207 |
| CepurR40.VG026700 | 0.005 | 0.028 | CepurGG1.UG154800 | 0.003 | 0.008 |
| CepurR40.VG027900 | 0.02 | 0.092 | CepurGG1.UG236700 | 0.009 | 0.072 |
| CepurR40.VG028400 | 0.02 | 0.065 | CepurGG1.UG146500 | 0.017 | 0.035 |
| CepurR40.VG030600 | 0.275 | 0.7 | CepurGG1.UG197100 | 0.122 | 0.357 |
| CepurR40.VG032800 | 0.015 | 0.104 | CepurGG1.UG307300 | 0.017 | 0.065 |
| CepurR40.VG033700 | 0.01 | 0.016 | CepurGG1.UG078100 | 0.002 | 0.025 |
| CepurR40.VG033900 | 0.011 | 0.24 | CepurGG1.UG218700 | 0.209 | 0.491 |
| CepurR40.VG034100 | 0.035 | 0.119 | CepurGG1.UG256900 | 0.037 | 0.091 |
| CepurR40.VG035600 | 0.053 | 0.12 | CepurGG1.UG027600 | 0.04 | 0.085 |

| | | | | | |
|---|---|---|---|---|---|
| CepurR40.VG036900 | 0.024 | 0.12 | CepurGG1.UG323900 | 0.013 | 0.115 |
| CepurR40.VG038300 | 0* | 0.193 | CepurGG1.UG056200 | 0.002 | 0.116 |
| CepurR40.VG038900 | 0.047 | 0.137 | CepurGG1.UG006700 | 0.03 | 0.062 |
| CepurR40.VG039900 | 0.002 | 0* | CepurGG1.UG006400 | 0.01 | 0.016 |
| CepurR40.VG043900 | 0.045 | 0.081 | CepurGG1.UG120100 | 0.024 | 0.074 |
| CepurR40.VG044000 | 0* | 0.036 | CepurGG1.UG119900 | 0.008 | 0* |
| CepurR40.VG044300 | 0.009 | 0.2 | CepurGG1.UG328100 | 0.009 | 0.117 |
| CepurR40.VG045300 | 0.011 | 0.131 | CepurGG1.UG198600 | 0.008 | 0.063 |
| CepurR40.VG047000 | 0* | 0.084 | CepurGG1.UG333100 | 0.17 | 0.346 |
| CepurR40.VG047100 | 0.001 | 0* | CepurGG1.UG034400 | 0.003 | 0.019 |
| CepurR40.VG051900 | 0.003 | 0.031 | CepurGG1.UG092100 | 0.01 | 0.003 |
| CepurR40.VG054000 | 0.006 | 0.006 | CepurGG1.UG082200 | 0.005 | 0* |
| CepurR40.VG054100 | 0* | 0.006 | CepurGG1.UG082100 | 0.001 | 0.007 |
| CepurR40.VG054200 | 0.014 | 0* | CepurGG1.UG082000 | 0.074 | 0.115 |
| CepurR40.VG054900 | 0.01 | 0.032 | CepurGG1.UG033700 | 0.004 | 0.002 |
| CepurR40.VG055100 | 0.009 | 0.096 | CepurGG1.UG033000 | 0.006 | 0.092 |
| CepurR40.VG055800 | 0.007 | 0.012 | CepurGG1.UG263600 | 0.057 | 0.077 |
| CepurR40.VG056000 | 0.011 | 0.025 | CepurGG1.UG263800 | 0.014 | 0* |
| CepurR40.VG060100 | 0.049 | 0.078 | CepurGG1.UG105200 | 0.026 | 0.082 |
| CepurR40.VG061900 | 0.001 | 0.019 | CepurGG1.UG045500 | 0.002 | 0.003 |
| CepurR40.VG064100 | 0.004 | 0.006 | CepurGG1.UG089800 | 0.001 | 0* |
| CepurR40.VG064200 | 0.003 | 0* | CepurGG1.UG089900 | 0.01 | 0.009 |
| CepurR40.VG064500 | 0.016 | 0.003 | CepurGG1.UG045800 | 0.006 | 0.013 |
| CepurR40.VG064700 | 0.004 | 0.007 | CepurGG1.UG045600 | 0.006 | 0.024 |
| CepurR40.VG065300 | 0.007 | 0.015 | CepurGG1.UG090300 | 0.003 | 0.004 |
| CepurR40.VG065400 | 0.004 | 0.011 | CepurGG1.UG090400 | 0.006 | 0.006 |
| CepurR40.VG065700 | 0.005 | 0.006 | CepurGG1.UG094900 | 0.032 | 0.055 |
| CepurR40.VG065900 | 0.013 | 0.005 | CepurGG1.UG086300 | 0.007 | 0.011 |
| CepurR40.VG066000 | 0.002 | 0.022 | CepurGG1.UG086200 | 0.002 | 0.009 |
| CepurR40.VG066300 | 0.007 | 0.014 | CepurGG1.UG084200 | 0.004 | 0.008 |
| CepurR40.VG067000 | 0.003 | 0.039 | CepurGG1.UG083500 | 0.005 | 0.019 |
| CepurR40.VG067600 | 0.005 | 0.039 | CepurGG1.UG086500 | 0.004 | 0.033 |
| CepurR40.VG067800 | 0.007 | 0.016 | CepurGG1.UG086600 | 0.003 | 0.01 |
| CepurR40.VG068100 | 0.009 | 0.015 | CepurGG1.UG086800 | 0.002 | 0* |
| CepurR40.VG068300 | 0.015 | 0.002 | CepurGG1.UG088000 | 0.004 | 0.014 |
| CepurR40.VG068700 | 0.01 | 0.089 | CepurGG1.UG098700 | 0.003 | 0.052 |
| CepurR40.VG069400 | 0.012 | 0.028 | CepurGG1.UG099600 | 0.013 | 0.021 |
| CepurR40.VG069500 | 0.01 | 0.021 | CepurGG1.UG099700 | 0.004 | 0.019 |
| CepurR40.VG069700 | 0.002 | 0.017 | CepurGG1.UG099900 | 0.002 | 0.007 |
| CepurR40.VG069900 | 0.007 | 0.013 | CepurGG1.UG100000 | 0.001 | 0.011 |
| CepurR40.VG070200 | 0.006 | 0.017 | CepurGG1.UG100800 | 0.005 | 0.04 |
| CepurR40.VG072900 | 0.042 | 0.111 | CepurGG1.UG323700 | 0.048 | 0.087 |
| CepurR40.VG073300 | 0.026 | 0.144 | CepurGG1.UG304700 | 0.033 | 0.14 |
| CepurR40.VG073500 | 0.022 | 0.075 | CepurGG1.UG149000 | 0.015 | 0.052 |
| CepurR40.VG074300 | 0.003 | 0.051 | CepurGG1.UG013700 | 0.003 | 0.014 |

| | | | | | |
|---|---|---|---|---|---|
| CepurR40.VG075100 | 0.033 | 0.112 | CepurGG1.UG228700 | 0.017 | 0.097 |
| CepurR40.VG075600 | 0.018 | 0.131 | CepurGG1.UG249900 | 0.011 | 0.109 |
| CepurR40.VG076700 | 0.013 | 0* | CepurGG1.UG275500 | 0.01 | 0.034 |
| CepurR40.VG078300 | 0.004 | 0* | CepurGG1.UG132500 | 0.005 | 0.013 |
| CepurR40.VG078400 | 0.01 | 0.012 | CepurGG1.UG162300 | 0.007 | 0.017 |
| CepurR40.VG078700 | 0.001 | 0* | CepurGG1.UG162900 | 0.009 | 0.042 |
| CepurR40.VG079400 | 0.051 | 0.195 | CepurGG1.UG163200 | 0.029 | 0.156 |
| CepurR40.VG079700 | 0.003 | 0.01 | CepurGG1.UG228400 | 0.006 | 0* |
| CepurR40.VG080000 | 0.002 | 0.041 | CepurGG1.UG228500 | 0.008 | 0.089 |
| CepurR40.VG080500 | 0.051 | 0.134 | CepurGG1.UG154200 | 0.028 | 0.112 |
| CepurR40.VG081000 | 0.007 | 0.078 | CepurGG1.UG321500 | 0.009 | 0.068 |
| CepurR40.VG081100 | 0.093 | 0.904 | CepurGG1.UG068400 | 0.082 | 0.422 |
| CepurR40.VG083300 | 0.027 | 0.133 | CepurGG1.UG147300 | 0.015 | 0.088 |
| CepurR40.VG085400 | 0.029 | 0.141 | CepurGG1.UG159500 | 0.013 | 0.109 |
| CepurR40.VG089900 | 0.026 | 0.043 | CepurGG1.UG304400 | 0.018 | 0.06 |
| CepurR40.VG090400 | 0.003 | 0* | CepurGG1.UG097100 | 0* | 0* |
| CepurR40.VG092200 | 0.06 | 0.16 | CepurGG1.UG204000 | 0.07 | 0.288 |
| CepurR40.VG092400 | 0.01 | 0.017 | CepurGG1.UG270100 | 0.012 | 0.036 |
| CepurR40.VG093200 | 0.017 | 0.098 | CepurGG1.UG320200 | 0.017 | 0.06 |
| CepurR40.VG094300 | 0.024 | 0.124 | CepurGG1.UG026500 | 0.02 | 0.043 |
| CepurR40.VG097000 | 0.019 | 0.042 | CepurGG1.UG270600 | 0.021 | 0.064 |
| CepurR40.VG097900 | 0.022 | 0.15 | CepurGG1.UG182000 | 0.016 | 0.11 |
| CepurR40.VG098100 | 0.042 | 0.171 | CepurGG1.UG294600 | 0.018 | 0.12 |
| CepurR40.VG099100 | 0.313 | 0.822 | CepurGG1.UG344800 | 0.096 | 0.11 |
| CepurR40.VG100600 | 0.016 | 0.091 | CepurGG1.UG344300 | 0.019 | 0.081 |
| CepurR40.VG104400 | 0.002 | 0.017 | CepurGG1.UG276000 | 0.003 | 0.006 |
| CepurR40.VG104500 | 0* | 0.028 | CepurGG1.UG276100 | 0.001 | 0.002 |
| CepurR40.VG104800 | 0.004 | 0.006 | CepurGG1.UG276200 | 0* | 0.006 |
| CepurR40.VG105700 | 0.007 | 0.026 | CepurGG1.UG167600 | 0.008 | 0.013 |
| CepurR40.VG105900 | 0.005 | 0.062 | CepurGG1.UG167200 | 0.002 | 0.022 |
| CepurR40.VG106100 | 0.001 | 0.029 | CepurGG1.UG168900 | 0.001 | 0.036 |
| CepurR40.VG106400 | 0.042 | 0.057 | CepurGG1.UG169300 | 0.008 | 0.006 |
| CepurR40.VG106600 | 0.015 | 0.024 | CepurGG1.UG166500 | 0.006 | 0.021 |
| CepurR40.VG107000 | 0.011 | 0.006 | CepurGG1.UG161800 | 0* | 0.007 |
| CepurR40.VG108600 | 0.007 | 0.011 | CepurGG1.UG054300 | 0.059 | 0.02 |
| CepurR40.VG109000 | 0.005 | 0.008 | CepurGG1.UG191400 | 0.007 | 0.025 |
| CepurR40.VG113600 | 0.007 | 0.055 | CepurGG1.UG054000 | 0.019 | 0* |
| CepurR40.VG115500 | 0.027 | 0.199 | CepurGG1.UG144400 | 0.051 | 0.218 |
| CepurR40.VG116100 | 0.04 | 0.128 | CepurGG1.UG008600 | 0.034 | 0.073 |
| CepurR40.VG117400 | 0.093 | 0.189 | CepurGG1.UG112300 | 0.044 | 0.128 |
| CepurR40.VG117900 | 0.018 | 0.087 | CepurGG1.UG298700 | 0.017 | 0.076 |
| CepurR40.VG118800 | 0.06 | 0.067 | CepurGG1.UG065400 | 0.041 | 0.005 |
| CepurR40.VG119200 | 0.015 | 0.105 | CepurGG1.UG307800 | 0.015 | 0.026 |
| CepurR40.VG120100 | 0.021 | 0.109 | CepurGG1.UG062700 | 0.015 | 0.113 |
| CepurR40.VG120800 | 0.004 | 0.033 | CepurGG1.UG258600 | 0.007 | 0.006 |
| CepurR40.VG122500 | 0.009 | 0.009 | CepurGG1.UG299800 | 0.011 | 0.017 |
| CepurR40.VG122800 | 0.003 | 0.006 | CepurGG1.UG283100 | 0.003 | 0.017 |

| | | | | | |
|---|---|---|---|---|---|
| CepurR40.VG126200 | 0.009 | 0.01 | CepurGG1.UG028800 | 0.005 | 0.023 |
| CepurR40.VG128400 | 0.008 | 0.144 | CepurGG1.UG182100 | 0* | 0.058 |
| CepurR40.VG128700 | 0.037 | 0.06 | CepurGG1.UG188000 | 0.025 | 0.072 |
| CepurR40.VG129500 | 0.046 | 0.156 | CepurGG1.UG040900 | 0.049 | 0.113 |
| CepurR40.VG130000 | 0.01 | 0.112 | CepurGG1.UG064500 | 0.006 | 0.081 |
| CepurR40.VG130100 | 0* | 0.116 | CepurGG1.UG319900 | 0* | 0.085 |
| CepurR40.VG131500 | 0.038 | 0.143 | CepurGG1.UG298400 | 0.054 | 0.2 |
| CepurR40.VG131900 | 0.004 | 0.059 | CepurGG1.UG325300 | 0.005 | 0.07 |
| CepurR40.VG133700 | 0.075 | 0.249 | CepurGG1.UG320800 | 0.044 | 0.28 |
| CepurR40.VG134300 | 0.026 | 0.133 | CepurGG1.UG224900 | 0.012 | 0.084 |
| CepurR40.VG136000 | 0.025 | 0.123 | CepurGG1.UG279000 | 0.005 | 0.092 |
| CepurR40.VG136300 | 0.014 | 0.268 | CepurGG1.UG041200 | 0.01 | 0.158 |
| CepurR40.VG138200 | 0.055 | 0.302 | CepurGG1.UG176300 | 0.069 | 0.13 |
| CepurR40.VG139700 | 0.028 | 0.091 | CepurGG1.UG279100 | 0.025 | 0.101 |
| CepurR40.VG143000 | 0.052 | 0.25 | CepurGG1.UG219400 | 0.06 | 0.25 |
| CepurR40.VG144200 | 0.024 | 0.101 | CepurGG1.UG295200 | 0.015 | 0.149 |
| CepurR40.VG145400 | 0.017 | 0.095 | CepurGG1.UG118400 | 0.019 | 0.118 |
| CepurR40.VG145800 | 0.003 | 0.084 | CepurGG1.UG340900 | 0.001 | 0.028 |
| CepurR40.VG151000 | 0.022 | 0.17 | CepurGG1.UG079300 | 0.011 | 0.103 |
| CepurR40.VG152200 | 0.016 | 0.152 | CepurGG1.UG223500 | 0.015 | 0.089 |
| CepurR40.VG153300 | 0.018 | 0.123 | CepurGG1.UG293900 | 0.008 | 0.141 |
| CepurR40.VG156200 | 0.013 | 0.087 | CepurGG1.UG293200 | 0.015 | 0.069 |
| CepurR40.VG156800 | 0.085 | 0.178 | CepurGG1.UG293100 | 0.036 | 0.126 |
| CepurR40.VG159200 | 0.019 | 0.162 | CepurGG1.UG294300 | 0.025 | 0.126 |
| CepurR40.VG160200 | 0.01 | 0.029 | CepurGG1.UG203700 | 0.007 | 0.015 |
| CepurR40.VG161200 | 0.023 | 0.135 | CepurGG1.UG278000 | 0.025 | 0.097 |
| CepurR40.VG161400 | 0.019 | 0.038 | CepurGG1.UG078400 | 0.025 | 0.032 |
| CepurR40.VG164200 | 0.037 | 0.106 | CepurGG1.UG009500 | 0.038 | 0.105 |
| CepurR40.VG164800 | 0.013 | 0.119 | CepurGG1.UG271000 | 0.002 | 0.135 |
| CepurR40.VG165000 | 0.025 | 0.078 | CepurGG1.UG282100 | 0.027 | 0.083 |
| CepurR40.VG168400 | 0.009 | 0.007 | CepurGG1.UG035400 | 0.005 | 0.02 |
| CepurR40.VG168700 | 0.008 | 0.019 | CepurGG1.UG035200 | 0* | 0.007 |
| CepurR40.VG169200 | 0.011 | 0.108 | CepurGG1.UG184400 | 0.001 | 0.099 |
| CepurR40.VG169800 | 0.026 | 0.121 | CepurGG1.UG326100 | 0.017 | 0.116 |
| CepurR40.VG170600 | 0.075 | 0.117 | CepurGG1.UG154000 | 0.041 | 0.078 |
| CepurR40.VG170700 | 0.003 | 0* | CepurGG1.UG240900 | 0.01 | 0.017 |
| CepurR40.VG171500 | 0.009 | 0.116 | CepurGG1.UG240700 | 0.014 | 0.072 |
| CepurR40.VG171900 | 0.068 | 0.135 | CepurGG1.UG066500 | 0.043 | 0.113 |
| CepurR40.VG173800 | 0 | 0* | CepurGG1.UG281900 | 0 | 0* |
| CepurR40.VG173900 | 0.01 | 0.083 | CepurGG1.UG126900 | 0.01 | 0.104 |
| CepurR40.VG174000 | 0.055 | 0.365 | CepurGG1.UG262300 | 0.025 | 0.22 |
| CepurR40.VG178500 | 0.012 | 0.038 | CepurGG1.UG321800 | 0.004 | 0.028 |
| CepurR40.VG179600 | 0.026 | 0.11 | CepurGG1.UG193500 | 0.015 | 0.077 |
| CepurR40.VG180100 | 0.012 | 0.186 | CepurGG1.UG296400 | 0.01 | 0.168 |
| CepurR40.VG180200 | 0.019 | 0.161 | CepurGG1.UG010600 | 0.009 | 0.12 |
| CepurR40.VG180900 | 0* | 0.017 | CepurGG1.UG102200 | 0 | 0.004 |
| CepurR40.VG181000 | 1.162 | 52.163 | CepurGG1.UG102300 | 0.044 | 0.136 |

| | | | | | |
|---|---|---|---|---|---|
| CepurR40.VG184300 | 0.031 | 0.143 | CepurGG1.UG024900 | 0.023 | 0.13 |
| CepurR40.VG185400 | 0.006 | 0.064 | CepurGG1.UG055500 | 0.022 | 0.064 |
| CepurR40.VG185800 | 0.047 | 0.155 | CepurGG1.UG288500 | 0.031 | 0.136 |
| CepurR40.VG188200 | 0.049 | 0.081 | CepurGG1.UG192800 | 0.026 | 0.067 |
| CepurR40.VG188400 | 0.024 | 0.085 | CepurGG1.UG235800 | 0.018 | 0.137 |
| CepurR40.VG189700 | 0.014 | 0.117 | CepurGG1.UG024400 | 0.035 | 0.084 |
| CepurR40.VG191500 | 0.025 | 0.09 | CepurGG1.UG182800 | 0.017 | 0.078 |
| CepurR40.VG193000 | 0.004 | 0.094 | CepurGG1.UG239700 | 0.006 | 0.107 |
| CepurR40.VG194900 | 0.007 | 0.018 | CepurGG1.UG173200 | 0.003 | 0.009 |
| CepurR40.VG198400 | 0.01 | 0.095 | CepurGG1.UG240400 | 0.007 | 0.084 |
| CepurR40.VG199100 | 0.008 | 0.077 | CepurGG1.UG053700 | 0.002 | 0.061 |
| CepurR40.VG200500 | 0.02 | 0.096 | CepurGG1.UG231500 | 0.012 | 0.078 |
| CepurR40.VG202500 | 0.006 | 0.017 | CepurGG1.UG137800 | 0.003 | 0.01 |
| CepurR40.VG202900 | 0.002 | 0.004 | CepurGG1.UG138200 | 0.002 | 0.009 |
| CepurR40.VG203100 | 0.004 | 0.071 | CepurGG1.UG138700 | 0.011 | 0.072 |
| CepurR40.VG203600 | 0.009 | 0.03 | CepurGG1.UG139400 | 0.012 | 0.04 |
| CepurR40.VG204500 | 0.016 | 0.019 | CepurGG1.UG141000 | 0.01 | 0.028 |
| CepurR40.VG204700 | 0.008 | 0.013 | CepurGG1.UG141200 | 0.018 | 0* |
| CepurR40.VG204800 | 0.051 | 0.116 | CepurGG1.UG141900 | 0.06 | 0.166 |
| CepurR40.VG204900 | 0.001 | 0.007 | CepurGG1.UG142100 | 0.003 | 0.008 |
| CepurR40.VG205000 | 0.018 | 0.177 | CepurGG1.UG142200 | 0.014 | 0.018 |
| CepurR40.VG205800 | 0.006 | 0* | CepurGG1.UG142400 | 0.008 | 0.044 |
| CepurR40.VG206500 | 0.007 | 0* | CepurGG1.UG189200 | 0.004 | 0.013 |
| CepurR40.VG206600 | 0.013 | 0.202 | CepurGG1.UG189300 | 0.016 | 0.179 |
| CepurR40.VG208800 | 0.009 | 0.027 | CepurGG1.UG014200 | 0.005 | 0.01 |
| CepurR40.VG209200 | 0* | 0.023 | CepurGG1.UG254700 | 0* | 0.022 |
| CepurR40.VG209400 | 0.012 | 0.029 | CepurGG1.UG059500 | 0.005 | 0.011 |
| CepurR40.VG210700 | 0.054 | 0.117 | CepurGG1.UG014000 | 0.059 | 0.089 |
| CepurR40.VG212700 | 0.066 | 0.102 | CepurGG1.UG106600 | 0.035 | 0.046 |
| CepurR40.VG213300 | 0.027 | 0.065 | CepurGG1.UG243100 | 0.014 | 0.107 |
| CepurR40.VG214400 | 0.02 | 0.085 | CepurGG1.UG196800 | 0.016 | 0.109 |
| CepurR40.VG215500 | 0.032 | 0.127 | CepurGG1.UG307500 | 0.012 | 0.092 |
| CepurR40.VG216600 | 0.004 | 0.071 | CepurGG1.UG285100 | 0.004 | 0.069 |
| CepurR40.VG217300 | 0.054 | 0.13 | CepurGG1.UG264500 | 0.033 | 0.091 |
| CepurR40.VG220400 | 0.007 | 0.151 | CepurGG1.UG058800 | 0.008 | 0.116 |
| CepurR40.VG220600 | 0.024 | 0.059 | CepurGG1.UG152900 | 0.019 | 0.107 |
| CepurR40.VG221200 | 0.021 | 0.117 | CepurGG1.UG220000 | 0.021 | 0.127 |
| CepurR40.VG221400 | 0.007 | 0.067 | CepurGG1.UG057600 | 0.011 | 0.046 |
| CepurR40.VG222100 | 0.02 | 0.147 | CepurGG1.UG249100 | 0.027 | 0.099 |
| CepurR40.VG226300 | 0* | 0.016 | CepurGG1.UG060600 | 0.007 | 0.011 |
| CepurR40.VG226700 | 0.049 | 0.247 | CepurGG1.UG064900 | 0.028 | 0.16 |
| CepurR40.VG227300 | 0.07 | 0.183 | CepurGG1.UG017500 | 0.006 | 0.105 |
| CepurR40.VG227600 | 0* | 0* | CepurGG1.UG111500 | 0.003 | 0.011 |
| CepurR40.VG233000 | 0.018 | 0.103 | CepurGG1.UG296500 | 0.01 | 0.088 |
| CepurR40.VG233900 | 0.02 | 0.185 | CepurGG1.UG025900 | 0.011 | 0.149 |
| CepurR40.VG234500 | 0.023 | 0.045 | CepurGG1.UG299400 | 0.017 | 0.073 |
| CepurR40.VG235300 | 0.098 | 0.294 | CepurGG1.UG071900 | 0.099 | 0.127 |

| | | | | | |
|---|---|---|---|---|---|
| CepurR40.VG235600 | 0.019 | 0.117 | CepurGG1.UG177100 | 0.016 | 0.045 |
| CepurR40.VG236000 | 0.038 | 0.351 | CepurGG1.UG069200 | 0.018 | 0.225 |
| CepurR40.VG236700 | 0.014 | 0.007 | CepurGG1.UG076500 | 0.006 | 0.023 |
| CepurR40.VG237800 | 0* | 0.01 | CepurGG1.UG317900 | 0.004 | 0* |
| CepurR40.VG242700 | 0 | 0* | CepurGG1.UG224200 | 0.003 | 0.009 |
| CepurR40.VG244200 | 0.018 | 0.14 | CepurGG1.UG125400 | 0.016 | 0.119 |
| CepurR40.VG245000 | 0.01 | 0.087 | CepurGG1.UG143200 | 0.003 | 0.068 |
| CepurR40.VG246200 | 0.018 | 0.064 | CepurGG1.UG067900 | 0.022 | 0.069 |
| CepurR40.VG247300 | 0.018 | 0.094 | CepurGG1.UG272600 | 0.02 | 0.071 |
| CepurR40.VG249700 | 0.012 | 0.022 | CepurGG1.UG185100 | 0.003 | 0.022 |
| CepurR40.VG250300 | 0.004 | 0.015 | CepurGG1.UG184900 | 0.004 | 0.018 |
| CepurR40.VG250400 | 0.004 | 0.018 | CepurGG1.UG155900 | 0* | 0.018 |
| CepurR40.VG250600 | 0.389 | 0.684 | CepurGG1.UG015400 | 0.043 | 0.068 |
| CepurR40.VG250800 | 0.005 | 0.02 | CepurGG1.UG015000 | 0.001 | 0* |
| CepurR40.VG251700 | 0.04 | 0.251 | CepurGG1.UG107100 | 0.033 | 0.298 |
| CepurR40.VG253000 | 0.02 | 0.132 | CepurGG1.UG339600 | 0.032 | 0.097 |
| CepurR40.VG253400 | 0.005 | 0* | CepurGG1.UG128100 | 0.005 | 0.011 |
| CepurR40.VG255400 | 0.018 | 0.128 | CepurGG1.UG209500 | 0.026 | 0.159 |
| CepurR40.VG257500 | 0.034 | 0.096 | CepurGG1.UG309400 | 0.027 | 0.075 |
| CepurR40.VG259000 | 0.01 | 0.14 | CepurGG1.UG210600 | 0.008 | 0.113 |
| CepurR40.VG259800 | 0* | 0.031 | CepurGG1.UG287500 | 0.003 | 0.026 |
| CepurR40.VG262500 | 0.019 | 0.034 | CepurGG1.UG261400 | 0.017 | 0.022 |
| CepurR40.VG262600 | 0.038 | 0.159 | CepurGG1.UG043700 | 0.027 | 0.127 |
| CepurR40.VG264300 | 0.009 | 0.029 | CepurGG1.UG047700 | 0.002 | 0.015 |
| CepurR40.VG264500 | 0.111 | 0.256 | CepurGG1.UG050100 | 0.067 | 0.141 |
| CepurR40.VG265400 | 0.038 | 0.174 | CepurGG1.UG212300 | 0.004 | 0.083 |
| CepurR40.VG267200 | 0.004 | 0.093 | CepurGG1.UG322300 | 0.003 | 0.122 |
| CepurR40.VG269700 | 0.013 | 0.109 | CepurGG1.UG260300 | 0.007 | 0.111 |
| CepurR40.VG270400 | 0.022 | 0.219 | CepurGG1.UG201900 | 0.014 | 0.147 |
| CepurR40.VG272300 | 0.014 | 0.059 | CepurGG1.UG341200 | 0.014 | 0.073 |
| CepurR40.VG273100 | 0.027 | 0.121 | CepurGG1.UG210200 | 0.03 | 0.099 |
| CepurR40.VG274500 | 0.011 | 0.032 | CepurGG1.UG180800 | 0.01 | 0* |
| CepurR40.VG274700 | 0.361 | 0.548 | CepurGG1.UG180500 | 0.009 | 0.014 |
| CepurR40.VG275500 | 0.005 | 0.008 | CepurGG1.UG269300 | 0.007 | 0.005 |
| CepurR40.VG275800 | 0.006 | 0* | CepurGG1.UG269600 | 0.005 | 0.018 |
| CepurR40.VG276900 | 0.002 | 0.016 | CepurGG1.UG268000 | 0.002 | 0.009 |
| CepurR40.VG277000 | 0.007 | 0.01 | CepurGG1.UG267800 | 0.013 | 0* |
| CepurR40.VG278100 | 0.006 | 0.012 | CepurGG1.UG267000 | 0.006 | 0.012 |
| CepurR40.VG280200 | 0.005 | 0.017 | CepurGG1.UG230200 | 0.002 | 0.019 |
| CepurR40.VG281200 | 0.02 | 0.104 | CepurGG1.UG342700 | 0.017 | 0.084 |
| CepurR40.VG282500 | 0.008 | 0.016 | CepurGG1.UG343500 | 0.005 | 0.01 |
| CepurR40.VG282700 | 0.007 | 0.037 | CepurGG1.UG227900 | 0.003 | 0.021 |
| CepurR40.VG282800 | 0* | 0.007 | CepurGG1.UG330600 | 0.008 | 0.013 |
| CepurR40.VG284000 | 0.032 | 0.124 | CepurGG1.UG260900 | 0.022 | 0.111 |
| CepurR40.VG284500 | 0.05 | 0.108 | CepurGG1.UG243900 | 0.04 | 0.061 |
| CepurR40.VG285600 | 0.046 | 0.135 | CepurGG1.UG131100 | 0.021 | 0.102 |
| CepurR40.VG287800 | 0.042 | 0.138 | CepurGG1.UG152400 | 0.035 | 0.149 |

| | | | | | |
|---|---|---|---|---|---|
| CepurR40.VG289600 | 0.06 | 0.064 | CepurGG1.UG159600 | 0.048 | 0.116 |
| CepurR40.VG290100 | 0.004 | 0.013 | CepurGG1.UG134700 | 0.007 | 0.024 |
| CepurR40.VG290200 | 0.004 | 0.009 | CepurGG1.UG134600 | 0.005 | 0* |
| CepurR40.VG290400 | 0.004 | 0.008 | CepurGG1.UG134400 | 0.004 | 0.001 |
| CepurR40.VG291700 | 0.003 | 0.014 | CepurGG1.UG134300 | 0.003 | 0.005 |
| CepurR40.VG291800 | 0.005 | 0.008 | CepurGG1.UG134100 | 0* | 0.008 |
| CepurR40.VG293000 | 0.002 | 0.021 | CepurGG1.UG133100 | 0.002 | 0.012 |
| CepurR40.VG293100 | 0* | 0.03 | CepurGG1.UG133200 | 0.006 | 0.005 |
| CepurR40.VG294700 | 0.003 | 0.057 | CepurGG1.UG275200 | 0* | 0.014 |
| CepurR40.VG295200 | 0* | 0.015 | CepurGG1.UG274700 | 0* | 0.006 |
| CepurR40.VG295500 | 0.003 | 0* | CepurGG1.UG274800 | 0* | 0.016 |
| CepurR40.VG296300 | 0.003 | 0.07 | CepurGG1.UG011200 | 0.003 | 0.068 |
| CepurR40.VG297000 | 0.017 | 0.074 | CepurGG1.UG095000 | 0.021 | 0.126 |
| CepurR40.VG297900 | 0.009 | 0.022 | CepurGG1.UG255700 | 0.002 | 0.006 |
| CepurR40.VG298800 | 0.002 | 0.017 | CepurGG1.UG171000 | 0.001 | 0.021 |
| CepurR40.VG299200 | 0.008 | 0.032 | CepurGG1.UG013600 | 0.005 | 0.028 |
| CepurR40.VG299600 | 0.048 | 0.099 | CepurGG1.UG215800 | 0.049 | 0.069 |
| CepurR40.VG300700 | 0.038 | 0.058 | CepurGG1.UG066300 | 0.059 | 0.121 |
| CepurR40.VG301200 | 0.118 | 0.23 | CepurGG1.UG285500 | 0.046 | 0.095 |
| CepurR40.VG301500 | 0.092 | 0.531 | CepurGG1.UG027500 | 0.026 | 0.186 |
| CepurR40.VG302900 | 0.03 | 0.121 | CepurGG1.UG152100 | 0.022 | 0.053 |
| CepurR40.VG304200 | 0.011 | 0.103 | CepurGG1.UG055400 | 0.009 | 0.064 |
| CepurR40.VG306500 | 0.016 | 0.093 | CepurGG1.UG218100 | 0.018 | 0.067 |
| CepurR40.VG307400 | 0.069 | 0.225 | CepurGG1.UG185700 | 0.04 | 0.16 |
| CepurR40.VG309000 | 0.088 | 0.25 | CepurGG1.UG281500 | 0.029 | 0.138 |
| CepurR40.VG310800 | 0.027 | 0.161 | CepurGG1.UG145500 | 0.014 | 0.153 |
| CepurR40.VG310900 | 0.043 | 0.186 | CepurGG1.UG284400 | 0.049 | 0.061 |
| CepurR40.VG311800 | 0.006 | 0.006 | CepurGG1.UG075500 | 0.005 | 0.007 |
| CepurR40.VG312500 | 0.001 | 0.01 | CepurGG1.UG032000 | 0.002 | 0.007 |
| CepurR40.VG313300 | 0.014 | 0.084 | CepurGG1.UG210800 | 0.021 | 0.086 |
| CepurR40.VG313700 | 0.003 | 0.014 | CepurGG1.UG178200 | 0.001 | 0* |
| CepurR40.VG315200 | 0.01 | 0.025 | CepurGG1.UG035000 | 0* | 0.014 |
| CepurR40.VG324400 | 0.012 | 0.053 | CepurGG1.UG122500 | 0.011 | 0.048 |
| CepurR40.VG325500 | 0.042 | 0.088 | CepurGG1.UG345000 | 0.038 | 0.092 |
| CepurR40.VG326300 | 0.004 | 0.11 | CepurGG1.UG186300 | 0.002 | 0.097 |
| CepurR40.VG329200 | 0.007 | 0.026 | CepurGG1.UG174100 | 0* | 0.005 |
| CepurR40.VG329600 | 0.004 | 0.012 | CepurGG1.UG174300 | 0.004 | 0.005 |
| CepurR40.VG329800 | 0.01 | 0.023 | CepurGG1.UG174500 | 0.005 | 0.005 |
| CepurR40.VG329900 | 0.204 | 0.184 | CepurGG1.UG096200 | 0.019 | 0.041 |
| CepurR40.VG331000 | 0.012 | 0.019 | CepurGG1.UG331000 | 0* | 0.013 |
| CepurR40.VG331200 | 0.008 | 0.012 | CepurGG1.UG330800 | 0.004 | 0.013 |
| CepurR40.VG331300 | 0.01 | 0.018 | CepurGG1.UG330700 | 0.009 | 0.013 |
| CepurR40.VG331600 | 0.009 | 0.022 | CepurGG1.UG336200 | 0.006 | 0.02 |
| CepurR40.VG331800 | 0.07 | 0.041 | CepurGG1.UG003100 | 0.034 | 0.025 |
| CepurR40.VG331900 | 0.001 | 0.005 | CepurGG1.UG003300 | 0 | 0.004 |
| CepurR40.VG332600 | 0.002 | 0.017 | CepurGG1.UG002400 | 0.009 | 0.01 |
| CepurR40.VG332700 | 0.004 | 0.015 | CepurGG1.UG002600 | 0.008 | 0.005 |

| | | | | | |
|---|---|---|---|---|---|
| CepurR40.VG333200 | 0.043 | 0.25 | CepurGG1.UG289100 | 0.041 | 0.239 |
| CepurR40.VG334400 | 0.002 | 0.01 | CepurGG1.UG263200 | 0.004 | 0.01 |
| CepurR40.VG335300 | 0.015 | 0.011 | CepurGG1.UG335300 | 0.008 | 0.017 |
| CepurR40.VG335400 | 0* | 0.018 | CepurGG1.UG335200 | 0.007 | 0.007 |
| CepurR40.VG335800 | 0.003 | 0.017 | CepurGG1.UG335500 | 0.002 | 0.006 |
| CepurR40.VG336000 | 0.004 | 0.043 | CepurGG1.UG335600 | 0* | 0.018 |
| CepurR40.VG336200 | 0.003 | 0.008 | CepurGG1.UG335800 | 0.003 | 0.011 |
| CepurR40.VG336600 | 0.005 | 0.044 | CepurGG1.UG077200 | 0.005 | 0.022 |
| CepurR40.VG337300 | 0.003 | 0.021 | CepurGG1.UG158900 | 0.004 | 0.016 |
| CepurR40.VG337700 | 0.005 | 0.023 | CepurGG1.UG158800 | 0.002 | 0.005 |
| CepurR40.VG338000 | 0.013 | 0.008 | CepurGG1.UG158700 | 0.016 | 0.012 |
| CepurR40.VG338100 | 0.042 | 0.113 | CepurGG1.UG158600 | 0.027 | 0.176 |
| CepurR40.VG338200 | 0.011 | 0.007 | CepurGG1.UG158500 | 0.011 | 0.004 |
| CepurR40.VG338800 | 0.013 | 0.062 | CepurGG1.UG157600 | 0* | 0 |
| CepurR40.VG339100 | 0.006 | 0.016 | CepurGG1.UG157700 | 0.004 | 0.003 |
| CepurR40.VG339300 | 0.004 | 0.028 | CepurGG1.UG157900 | 0.003 | 0.047 |
| CepurR40.VG339400 | 0.002 | 0.098 | CepurGG1.UG158000 | 0.007 | 0.045 |

### Supplemental Figures

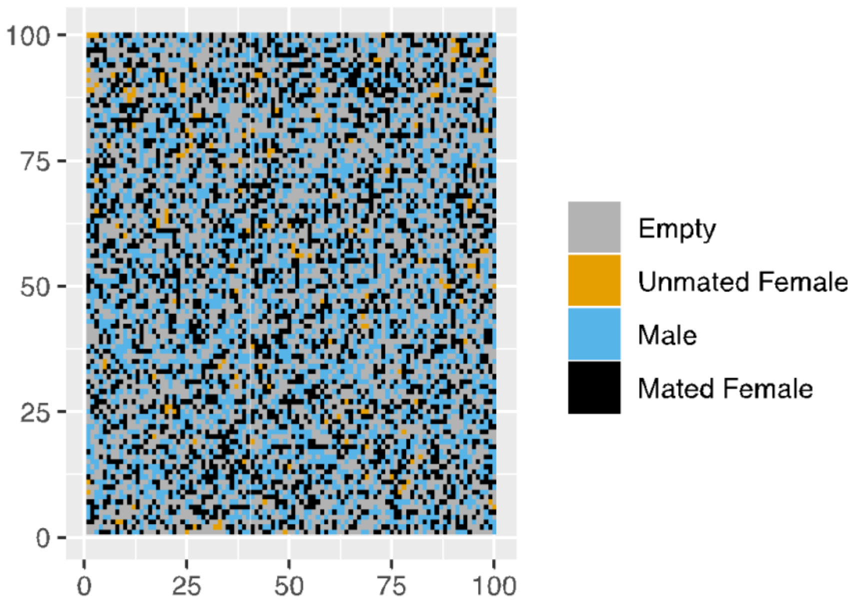


**Figure S1. Example of the population at the end of one run of the simulation.** Each cell is a patch that could or could not hold an individual. In the above example, the population size was 6000 and the sex ratio was one to one.
